# Combinatorial base editing couples disease correction with lineage amplification in hematopoietic stem and progenitor cells

**DOI:** 10.64898/2026.04.13.718029

**Authors:** Kun Jia, Eric Soupene, Roshani Sinha, Benjamin J. Lesch, Michael A. Pendergast, Rachel Choi, Xinyi Zhang, Elisabetta M. Foppiani, Zachary Kostamo, Simon N. Chu, Devesh Sharma, Xinjie Yu, Marco Cordero, Mark C. Walters, Tippi C. MacKenzie, Vivien A. Sheehan, M. Kyle Cromer

## Abstract

First-generation genome editing therapies have largely focused on correcting or compensating for pathogenic variants. However, as these approaches enter the clinic, emerging biological constraints limit maximal therapeutic impact. Because globin genes are activated late during erythroid differentiation, genome-corrected hematopoietic stem and progenitor cells (HSPCs) gain little selective advantage in the bone marrow. Here, we establish a strategy that links therapeutic genome edits to an erythroid fitness-enhancing allele to amplify the output of clinically relevant cells. We develop a multiplex base editing strategy that couples fetal hemoglobin (HbF) reactivation with erythroid lineage expansion. Introduction of a naturally occurring erythropoietin receptor truncation (*tEPOR*) associated with benign erythrocytosis increased erythroid cell production without impairing viability or differentiation. Combinatorial editing of *tEPOR* together with the *BCL11A* erythroid enhancer and *HBG1/2* promoters in healthy donor, sickle cell disease, and β-thalassemia HSPCs synergistically increased erythroid proliferation and HbF expression beyond single base-edited or Casgevy-treated controls. Multiplex base-edited HSPCs retained long-term lineage repopulation and engraftment capacity *in vivo*, establishing a modular strategy that pairs disease correction with lineage amplification to improve therapeutic potency.

## Introduction

Sickle cell disease (SCD) and β-thalassemia were among the first disorders understood at the molecular level and have long served as model indications for genetic medicine. Decades of work in globin biology recently culminated in the FDA approval of the first CRISPR-based therapy, Casgevy (exa-cel), which reactivates fetal hemoglobin (HbF; a tetramer comprised of two gamma (γ)- and two alpha (α)-globin chains) through disruption of the *BCL11A* erythroid enhancer^1^. While precise genome editing fulfills a long-standing goal in therapeutic science, its clinical implementation has revealed biological bottlenecks that limit maximal therapeutic impact.

One key limitation arises from the developmental timing of globin expression, which is activated late during erythroid maturation after hematopoietic stem and progenitor cells (HSPCs) have already undergone lineage commitment. Consequently, genome-edited HSPCs gain little intrinsic advantage in the bone marrow and are not biased toward producing the lineage most relevant for clinical benefit in hemoglobinopathies—the red blood cell (RBC). In the absence of a selective advantage, therapeutic efficacy depends on both the ability to mobilize sufficient numbers of HSPCs for editing, a known challenge in SCD^2^, and achieving high levels of edited HSPC engraftment. The latter is only possible through myeloablative conditioning which is associated with substantial toxicities including organ damage, infertility, prolonged immunosuppression, and increased risk of secondary malignancy^3,4^. Strategies that increase the erythroid contribution of edited cells could therefore reduce dependence on large input cell doses and reliance on intensive conditioning while improving therapeutic output.

In parallel, concerns have emerged regarding genome editing approaches that rely on Cas9-induced DNA double-strand breaks (DSBs), particularly in long-term repopulating HSCs^5^. DSB repair can generate unintended genomic alterations—including large deletions, translocations, and inversions^6,7^—and emerging evidence suggests that disruption of the *BCL11A* enhancer itself may impair erythroid differentiation and RBC output^8^. These limitations motivate the development of genome engineering strategies that enhance therapeutic efficacy while minimizing genotoxic risk and preserving erythropoiesis.

Base editors represent a newer class of CRISPR-derived tools capable of introducing precise nucleotide substitutions without programmed induction of DSBs or exogenous donor DNA templates^9^. These technologies have rapidly advanced from pre-clinical studies to early clinical trials^1,10–15,19,37^ (**Table S1**), underscoring their translational potential. Importantly, the predictable and scarless nature of base editing makes it particularly well suited for multiplex genome engineering^16,17^, enabling simultaneous installation of disease-corrective mutations together with protective or fitness-enhancing variants.

Here, we establish a framework for combinatorial base editing in primary human HSPCs that integrates disease correction with lineage amplification. We precisely introduce a naturally occurring single-nucleotide variant in the erythropoietin receptor (*EPOR*), originally identified in a Finnish Olympic cross-country skier with benign erythrocytosis, which truncates the receptor C-terminus and confers erythropoietin hypersensitivity^18,20,21^. We then combinatorially pair this allele (*tEPOR*) with HbF-inducing edits in the *BCL11A* erythroid enhancer^22^ and/or promoter region of the γ-globin locus (encoded by *HBG1* and *HBG2* genes)^19^, achieving efficient dual- and triple-editing at all three loci in healthy donor, SCD, and β-thalassemia patient-derived HSPCs. To our knowledge, prior studies have achieved at most two concurrent edits in primary HSPCs, often with variable efficiency; in contrast, we observe efficient installation of up to three edits across all targeted loci (**Table S2**). Addition of *tEPOR* amplifies erythroid output from HbF-activated cells, yielding substantial increases in total erythroid production and HbF levels while compensating for erythropoietic deficits associated with *BCL11A* enhancer disruption. Multiplex-edited HSPCs retained durable long-term engraftment potential in xenograft transplantation assays and outperformed Cas9-based approaches such as Casgevy, demonstrating robust human cell reconstitution despite the increased editing burden.

These findings illustrate how multiplex genome engineering can couple disease correction with functional enhancement to improve therapeutic performance. More broadly, demonstrating that primary HSPCs can tolerate and retain multiple cooperating edits expands the design space of genome-edited cell therapies, enabling integration of lineage amplification, immune evasion, antigen deletion, or epitope shielding within a single engineered product. Together, these results establish multiplex base editing as a modular and well-tolerated platform for engineering human HSPCs and provide a framework for next-generation stem cell therapies that combine corrective and fitness-enhancing edits to overcome emerging clinical bottlenecks.

## Results

### Base editing installs a natural EPOR variant that confers erythroid proliferative advantage

To enable precise installation of *EPOR* variants without generating DSBs, we designed cytosine base editor (CBE)-compatible single-guide RNAs (gRNAs) targeting exon 8 of *EPOR* to introduce premature stop codons upstream of the C-terminal SHP-1 binding motifs^23,24^. Two guides were selected: *EPOR*-sg1, which generates a nearby truncating variant, and *EPOR*-sg2, which recreates the exact naturally occurring mutation identified in the original erythrocytosis proband (**Figures 1A**). We first evaluated editing feasibility in HeLa cells by transfecting AncBE4max, an established CBE^25^, together with either *EPOR*-sg1 or *EPOR*-sg2 and collecting genomic DNA 72 hours later (**Figure S1B**). Sanger sequencing followed by EditR analysis^26^ confirmed efficient C-to-T editing at both target sites (**Figure S1A-B**), with each gRNA yielding ∼50-60% stop-codon alleles.

**Fig. 1:**
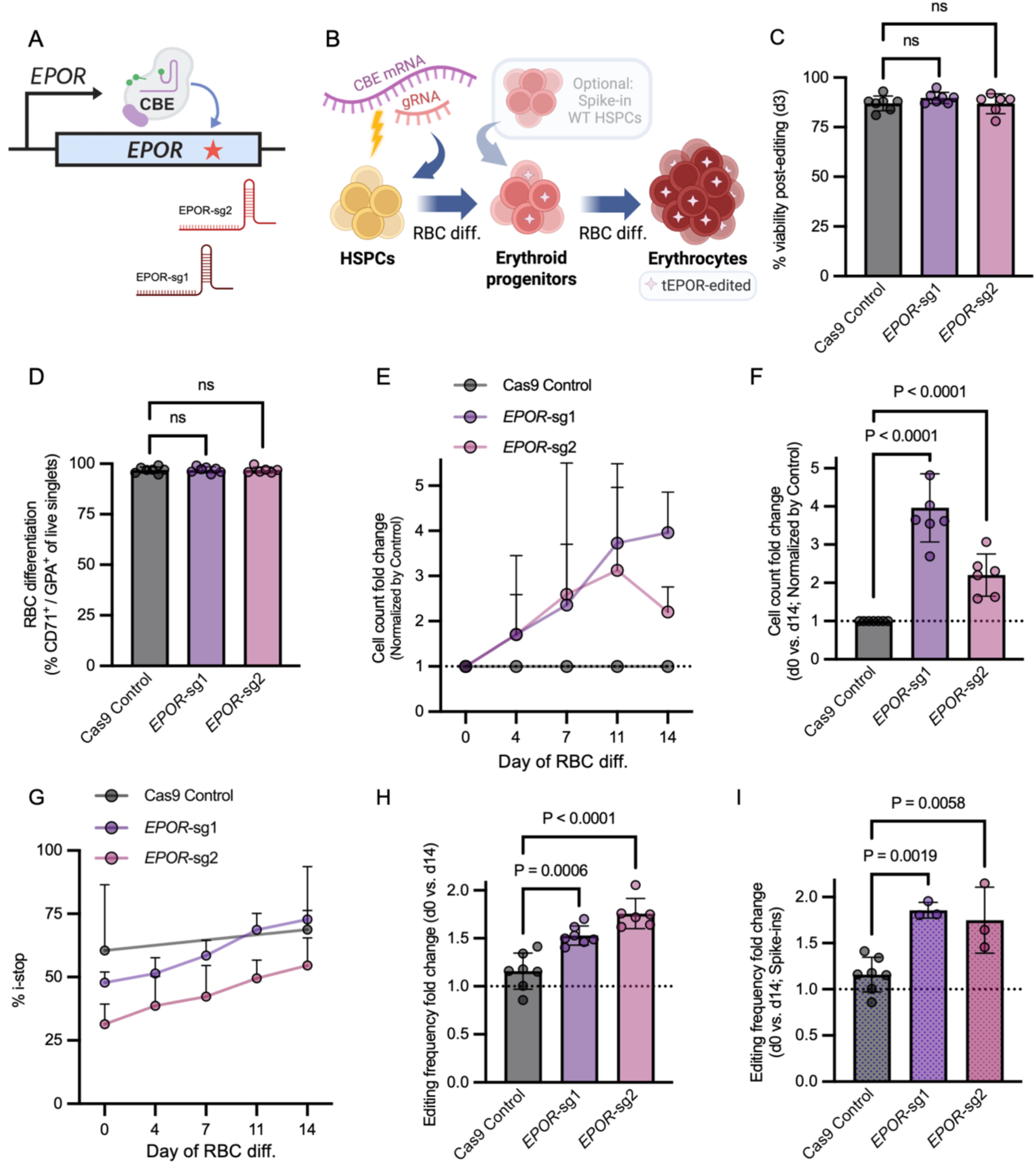
Precise *EPOR* truncation by base editing drives erythroid cell expansion. **A.** Schematic of the CBE editing strategy and gRNA sequences used to introduce the truncated *EPOR*. The protospacer adjacent motif (PAM) is shown in red, the protospacer in blue, predicted editing window in green. The “Olympic skier” variant is indicated for sg2. **B.** Schematic of the experimental workflow. Human CD34⁺ HSPCs were electroporated with CBE mRNA and gRNAs, and then subjected to *in vitro* erythroid differentiation. *EPOR*-edited cells are indicated by star symbols. In some experiments, wild-type (WT) HSPCs were “spiked-in” to model differentiation outcomes in a mixed population of edited and unedited cells. **C.** Percentage of viable cells at Day 3 post-editing measured by trypan blue exclusion assay across EPOR-edited conditions using CBE or control Cas9 edit at *CCR5* locus. **D.** *In vitro* erythroid differentiation frequency of edited HSPCs, assessed by flow cytometry as the percentage of CD71⁺/GPA⁺ cells among live singlets. **E.** Cell count fold change dynamics over the course of erythroid differentiation, normalized to control-edited condition at each timepoint. **F.** Total cell count fold change over course of erythroid differentiation, normalized to control. Statistical significance was assessed on non-normalized data using a linear mixed-effects model with donor as a random effect. *P*-values were adjusted using the Dunnett multiple comparisons method. **G.** Percentage of edited alleles at *tEPOR* allele in CD34⁺ HSPCs targeted with *EPOR*-sg1 or *EPOR*-sg2 over the course of erythroid differentiation (measured by NGS), or at the *CCR5* locus among control-edited HSPCs (measured by Sanger sequencing). **H-I.** Total fold enrichment of *tEPOR* editing frequencies over the course of erythroid differentiation among standard conditions (I) and with spike-in of unedited cells (J), both normalized to control. All columns represent mean values, and error bars indicate standard deviation (SD). Columns with dotted shading denote data from spike-in experiments. Unless otherwise specified, statistical significance was determined using one-way ANOVA with Dunnett’s multiple comparisons test (D-E, I-J) or linear mixed-effects models with donor as a random effect followed by Dunnett’s multiple comparisons test (G), comparing each *EPOR*-edited group to the control (*CCR5 Cas9*). ns, not significant (P > 0.05). Data are shown as individual measurements from independent donors (n = 7 total donors), with some donors represented more than once in selected conditions. Not all conditions were tested in all donors; therefore, sample availability varies across groups.

We next tested these strategies in human primary CD34⁺ HSPCs from mobilized healthy donors, electroporating cells with CBE mRNA and either *EPOR*-sg1 or *EPOR*-sg2 before initiating our established *in vitro* erythroid differentiation workflow^27^ (**Figure 1B** and **S2A**). Editing was well-tolerated, with comparable viability across all conditions (**Figure 1C**), and erythroid differentiation remained highly efficient across conditions when analyzed by flow cytometry at Day 14 of erythroid differentiation (**Figure 1D** and **S2B**). Cells were counted and genomic DNA was collected over the course of erythroid differentiation (Days 0, 4, 7, 11, and 14) to monitor proliferation dynamics of edited cell populations. In addition, we established both bulk-edited cultures and competitive co-cultures generated by mixing edited and unedited cells at a 1:1 ratio in order to model post-transplant competition between edited and unedited hematopoietic cells. We found that both CBE editing conditions conferred a proliferative advantage with *EPOR*-sg1 and *EPOR*-sg2 cultures achieving up to four-fold greater cell counts relative to control-treated cells (**Figure 1E**-**F** and **S2C**; *P* < 0.0001) with corresponding increases in edited cell frequency as determined by targeted next-generation sequencing (NGS) (**Figure 1G**-**H**; *P* ≤ 0.0006). These editing frequency fold changes were even more pronounced in spike-in cultures, indicating that *EPOR*-edited cells out-proliferated unedited cells over the course of erythroid differentiation (**Figure 1I**; *P* ≤ 0.0058).

To ensure a robust safety profile of these novel *EPOR* gRNAs, we then used *in silico* prediction tool COSMID^28,29^ to nominate candidate off-target editing sites throughout the genome. We found that all nominated sites had two or more mismatches in the protospacer sequence and that nearly all of these sites resided in non-coding intergenic or intronic regions of the genome (**Figure S3A**). Finally, transcriptomic data from prior work^30^ indicated that candidate off-target genes are minimally expressed throughout erythropoiesis, in contrast to the marked induction of *EPOR* during differentiation (**Figure S3B**). This suggests that even if off-target editing occurred, it would be unlikely to be biologically meaningful or to impact erythroid development.

### tEPOR multiplexing with HbF-reactivating edits drives selective erythroid expansion

Having established that CBE-mediated introduction of *tEPOR* enhances erythropoietic output, we next sought to pair this proliferative advantage with established therapeutic edits. Specifically, we replicated HbF-reactivating strategies that have shown clinical success, including base editing approaches that disrupt the erythroid enhancer of *BCL11A*^22^ or directly modify the *HBG* promoters^19^ to block repressive factor binding (**Figure 2A** and **Table S1**). The *BCL11A* strategy forms the basis of the first FDA-approved CRISPR therapy for sickle cell disease^11^, which restores HbF but provides no selective advantage to edited HSPCs because globin genes activate late in erythroid maturation. Therefore, we hypothesized that combining these therapeutic HbF-reactivating edits with *tEPOR* could enhance the output of HbF-expressing erythroid cells, thereby reducing the functional threshold of edited cell engraftment required for durable disease correction. However, the success of this concept will hinge on achieving high multiplex base editing rates at each intended target site in order to ensure that both HbF-reactivating edits and erythropoiesis-enhancing *tEPOR* edits occur in the same cell (**Figure 2B**).

**Fig. 2:**
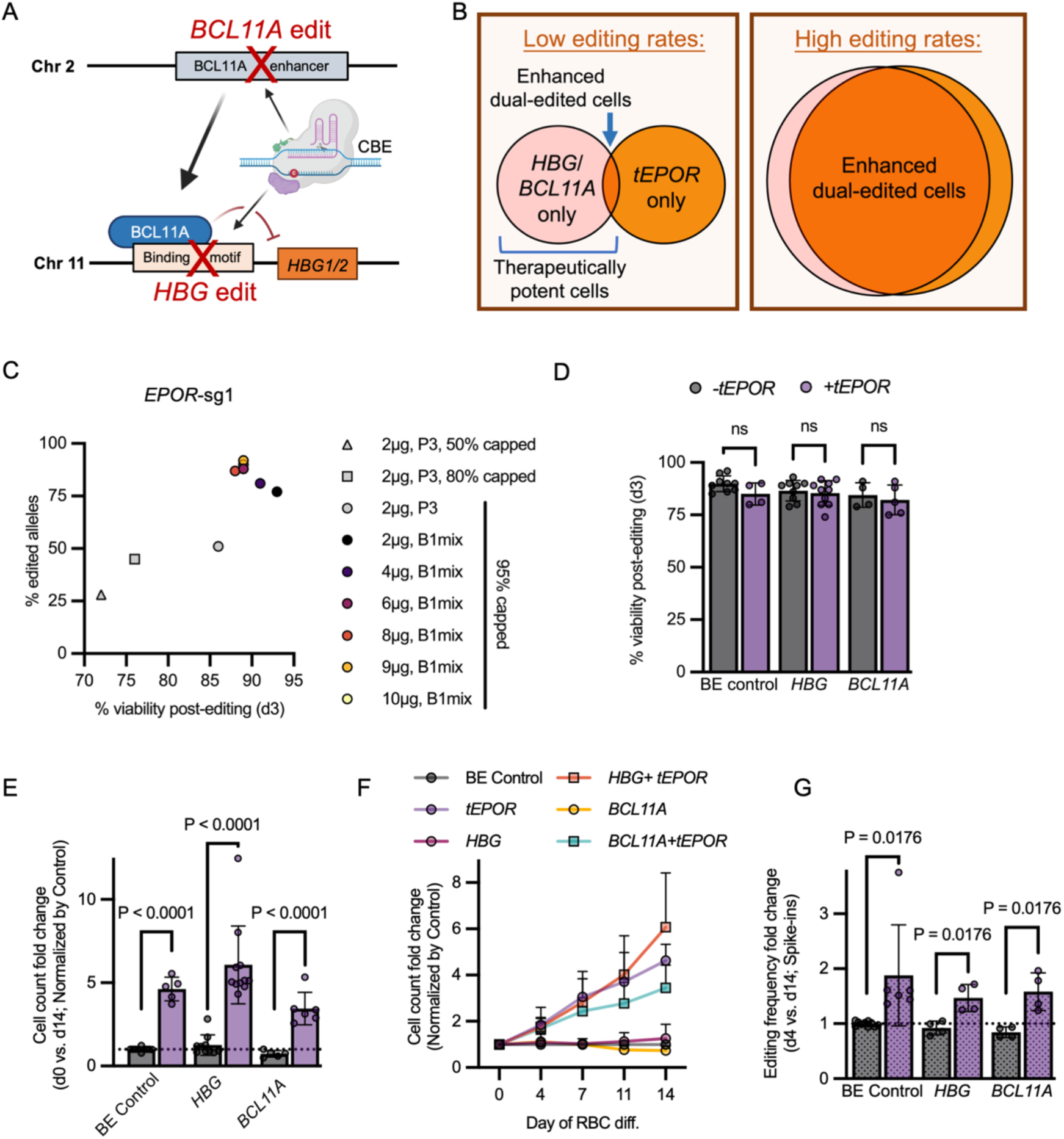
Multiplex base editing couples *EPOR* truncation with therapeutic *BCL11A* and *HBG* edits. **A.** Schematic of CBE-mediated fetal hemoglobin (HbF)-reactivation strategies targeting the *BCL11A* enhancer or BCL11A binding motif in *HBG1*/*2* promoters. **B.** Conceptual model of multiplex base editing outcomes when editing frequencies are low vs. high. **C.** Correlation between edited allele frequency and viability (quantified at Day 4 post-editing in human HSPCs) among optimized mRNA and electroporation conditions. **D.** Percentage of viable cells at Day 3 post-editing following single or multiplex base editing with CBE6. **E.** Total cell count fold change over course of erythroid differentiation, normalized to the BE control (*CCR5* CBE edit). Statistical significance was assessed on non-normalized data using a linear mixed-effects model with donor as a random effect. *P*-values were adjusted using the Holm method. **F.** Cell count fold change dynamics of edited cells over course of erythroid differentiation, normalized to control at each timepoint. **G.** Editing frequency fold change over course of erythroid differentiation with spike-in of unedited cells, normalized to control. Gray columns indicate samples without *tEPOR*, and purple columns depict samples with *tEPOR* co-editing. Columns depict mean ± SD. Columns with dotted shading indicate data from spike-in experiments. Unless otherwise indicated, statistical comparisons between +*tEPOR* and -*tEPOR* groups were performed using unpaired t-tests with Holm-Šidák correction for multiple comparisons (α = 0.05); ns, not significant (*P* > 0.05). Data are shown as individual measurements from independent donors (n = 7 total donors), with some donors represented more than once in selected conditions. Not all conditions were tested in all donors; therefore, sample availability varies across groups.

While our initial efforts demonstrated successful introduction of the intended *tEPOR* edit at nearly 50% of alleles (**Figure 1G**), we transitioned from AncBE4max to CBE6, a next-generation TadA-derived CBE engineered for high C-to-T conversion with markedly reduced A-to-G bystander activity, as the platform for subsequent optimization^31^. We then optimized CBE6 mRNA design and electroporation parameters to maximize base editing frequency in primary HSPCs. We first evaluated multiple biochemical features of the mRNA, including replacement of UTP with N1-methyl-pseudouridine during *in vitro* transcription, incorporation of CleanCap AG during capping, addition of RNase inhibitor, and the effect of using mRNA subjected to 48 or 72 hours of post-thaw storage prior to electroporation. These modifications—particularly N1m-U substitution, CleanCap AG capping (∼95% efficiency), water elution, and omission of RNase inhibitor—collectively yielded the highest levels of *EPOR*-sg1 editing (**Figure S4A**). Using this optimized mRNA configuration, we next performed dose-response testing (ranging 2-10 μg) across Lonza 4D-Nucleofector programs. This matrix identified CM-137 combined with 9 μg in B1mix buffer^35^ as the most effective condition, achieving up to 85% *EPOR* edited allele frequency while maintaining high cell viability (**Figure 2C** and **S4B**). Compared with the AncBE4max editor under non-optimized conditions, CBE6 substantially improved editing in CD34⁺ HSPCs, achieving editing efficiencies to a mean of 85.8% edited alleles at *BCL11A*, 74.8% at *HBG*, and 84.7% at *tEPOR*.

Using this optimized protocol, we deployed a multiplex CBE strategy that paired the proliferative *tEPOR* edit with each corrective edit individually, simultaneously targeting the *EPOR* premature-stop site and either the *BCL11A* intronic enhancer or the *HBG1/2* promoter motif in healthy donor primary HSPCs. This was achieved by electroporation of CBE6 mRNA along with gRNAs directed toward each intended target locus. In doing so, we found that dual-editing was well-tolerated, with high post-editing viability observed across control, single-, and dual-editing conditions (**Figure 2D**). We then tracked cell counts and editing frequencies over the course of erythroid differentiation. We observed a proliferative advantage (4-6-fold increase; *P* < 0.0001) among both the single *tEPOR* and dual *tEPOR*-edited conditions (**Figure 2E**-**F**). Similarly, we observed a consistent increase in edited cell frequency over the course of erythroid differentiation among all *tEPOR* single- and dual-edited conditions (**Figure 2F**; *P* = 0.0176). Interestingly, we observed a notable decrease in total cell count and editing frequencies by the end of erythroid differentiation among cells edited at *BCL11A* alone (**Figure 2E-F**), which reinforces the notion that ineffective erythropoiesis occurs when disrupting this master regulator gene^8^.

Together, these findings demonstrate that our optimized base editing protocol supports efficient dual-editing at independent loci and that combining clinical edits with *tEPOR* provides a robust proliferative advantage to edited cells without compromising viability or differentiation capacity.

### Triple-base editing drives erythroid expansion and enhances HbF induction

We next sought to assess whether our optimized protocol would allow efficient base editing of all three loci simultaneously. We hypothesized that combining *HBG*+*BCL11A* edits, to reactivate HbF, with *tEPOR*, to expand erythroid lineage, would maximize erythropoietic output of HbF-reactivated erythroid cells.

To monitor impact on HSPC viability and differentiation, we chose to benchmark single-, dual-, and triple-base editing against Casgevy (Cas9 editing at *BCL11A*) as well as the Cas9/AAV-editing platform (to integrate a full *HBB* ORF + *tEPOR* cDNA integration at the *HBA1* locus), which we previously developed for treatment of β-thalassemia^21,32^. We also performed Cas9 multiplex editing in an attempt to combine Casgevy with the *tEPOR* Cas9 edit defined previously. All of these strategies were deployed in CD34^+^ HSPCs and viability at Day 3 post-editing across all CBE treatments conditions was found to be not statistically different from unedited “Mock” cells, regardless of whether one, two, or three loci were targeted using base editors (**Figure 3A**). However, we observed a decrease in viability among both the Cas9 multiplex (Casgevy + *tEPOR*) and Cas9/AAV treatments (*P* ≤ 0.001), an effect that was eliminated in the latter condition when reducing the AAV dose (from 5,000 to 625 viral genomes per cell (MOI)), using manufactured AAV rather than prepped in-house, and adding AZD7648 to increase integration frequencies at lower MOI^33^. In spite of the observed cytotoxicity among dual Cas9- and Cas9/AAV editing conditions, all treatments supported robust erythroid maturation, generating high frequencies of CD71⁺/GPA⁺ cells at Day 14 of differentiation (**Figure 3B**).

**Fig. 3:**
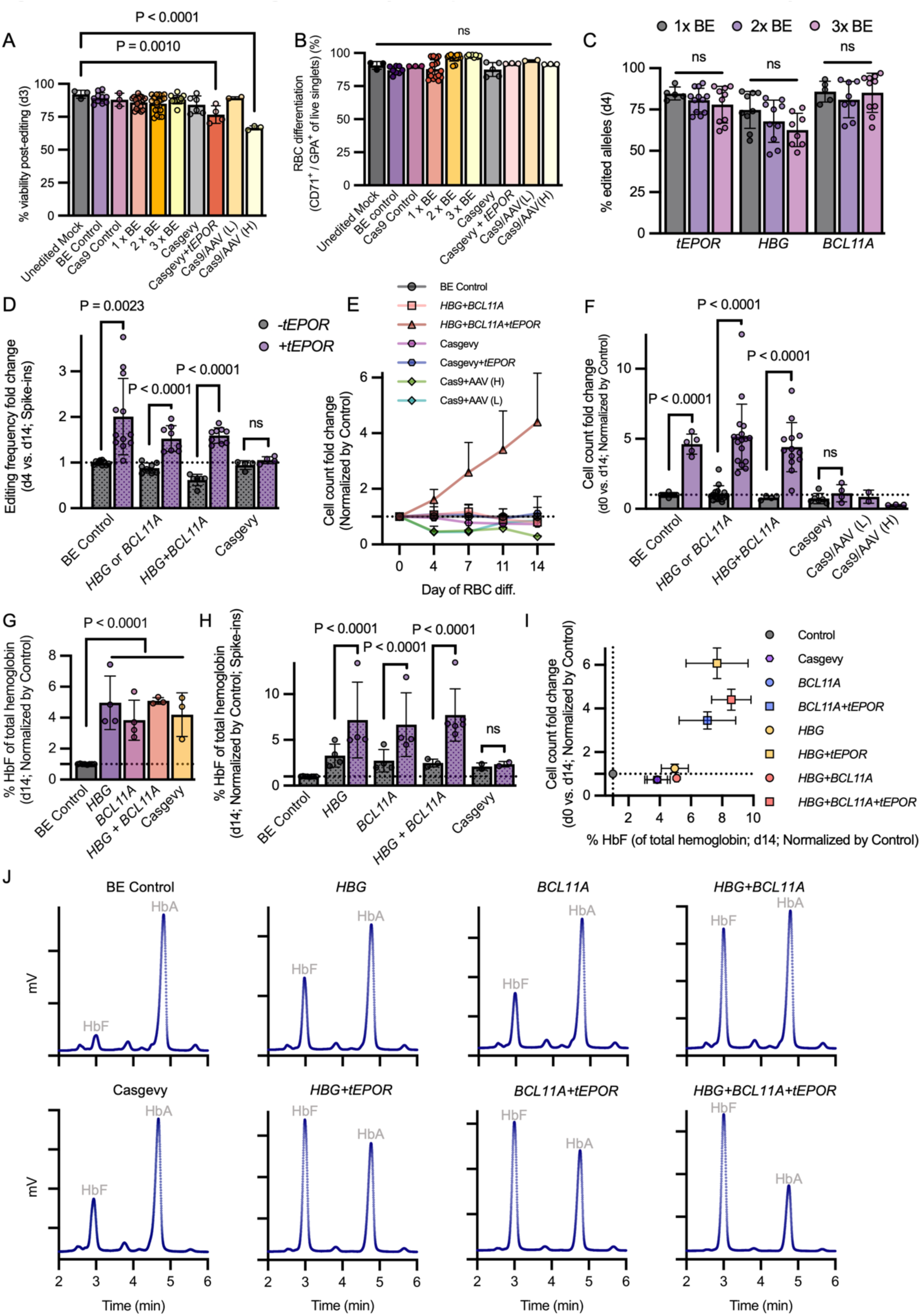
Combinatorial base editing enhances erythroid proliferation and HbF induction in healthy donor HSPCs. **A.** Percentage of viable cells at Day 3 post-editing across single- (1x), dual- (2x), and triple- (3x) BE conditions, Casgevy (*BCL11A* Cas9), Cas9/AAV (*HBB*-IRES-*tEPOR* into the *HBA1* locus; Cas9/AAV(L) = MOI 625 + AZD7648; Cas9/AAV(H) = MOI 5,000), BE control (*CCR5* CBE edit), Cas9 control (*CCR5* Cas9 edit), and mock (unedited control). Only statistically significant comparisons are annotated with P values; non-significant comparisons are not shown. **B.** Erythroid differentiation efficiency of edited HSPCs at end of erythroid differentiation. Data for BE control, 1x, and 2x BE conditions were reproduced and combined from Fig. 2F for comparison. **C.** Editing frequencies at Day 4 of erythroid differentiation at the *tEPOR, HBG,* and *BCL11A* loci across single-, dual-, and triple-editing conditions. **D.** Editing frequency fold change over course of erythroid differentiation with spike-in of unedited cells, normalized to control. Data for *HBG/BCL11A* conditions were reproduced from Fig. 2I for comparison. **E.** Cell count fold change dynamics over course of erythroid differentiation, normalized to control at each timepoint. Cas9/AAV condition integrated *HBB*-IRES-*tEPOR* at *HBA1* locus and AAV6 was delivered at high (H) or low (L) multiplicity of infection (MOI). **F.** Total cell count fold change over course of erythroid differentiation, normalized to control. Data for *HBG/BCL11A* conditions were reproduced from Fig. 2G. **G-H.** HbF (% of total hemoglobin) following erythroid differentiation, determined by hemoglobin tetramer HPLC and normalized to control. Total hemoglobin comprises HbF, HbA, and HbA2. Shown are single-edited conditions without spike-in (H) and dual-edited conditions with spike-in (I). **I.** Relationship between HbF induction and erythroid expansion across editing conditions. Error bars indicate SEM. **J.** Representative hemoglobin tetramer HPLC chromatograms from cell lysates. Peaks corresponding to fetal hemoglobin (HbF) and adult hemoglobin (HbA) are indicated. HbA2 was detected in all samples (eluting at ∼5.8 min), but is not labeled for clarity. Gray columns depict samples without *tEPOR*, and purple columns depict samples with *tEPOR* co-editing. Columns depict mean ± SD. Columns with dotted shading depict data from spike-in experiments. Statistical significance was determined using one-way ANOVA (A-C), unpaired t-tests (D) or linear mixed-effects models with donor as a random effect (F, G-H). *P*-values were adjusted for multiple comparisons (Holm or Dunnett). ns, not significant (*P* > 0.05). Data are shown as individual measurements from independent donors (n = 7 total donors), with some donors represented more than once in selected conditions. Not all conditions were tested in all donors; therefore, sample availability varies across groups.

Across all conditions, we isolated gDNA at Days 4 and 14 post-editing to monitor initial editing frequencies and to determine whether multiplexing with *tEPOR* imparts a proliferative advantage to edited cells over the course of erythroid differentiation. Although we observed a slight decrease in editing frequencies within 2x and 3x editing conditions compared to editing at a single target locus, these decreases were modest and did not achieve statistical significance (**Figure 3C**). Tracking editing dynamics from Day 4 to 14 revealed that all conditions receiving the *tEPOR* base edit showed increased editing frequencies (>50% in dual- and triple-edited conditions; **Figure 3D** and **S5A**; *P* ≤ 0.0023). As expected, we observed no increase in editing frequencies among control, *HBG-*, *BCL11A*-, and Casgevy-edited conditions.

To determine whether this enrichment of edited alleles translated into functional expansion, we next quantified total cell output following erythroid differentiation. Indeed, we observed a >4-fold increase in total cell counts among dual- and triple-base edited cells by the end of *in vitro* erythroid differentiation compared to dual *HBG+BCL11A*, Cas9/AAV, Casgevy, and Cas9 multiplex editing (**Figure3E-F**, and **S5B-C**; *P* < 0.0001). The lack of increased edited allele frequency or cell expansion when Casgevy was multiplexed with *tEPOR* indicates inefficient co-occurrence of the intended edits, potentially compounded by genotoxic stress associated with multiple DSBs. In addition, Cas9 targeting of *EPOR* generates heterogeneous indels^21^, only a subset of which encode functional *tEPOR* alleles, whereas BE precisely installs the desired allele. In contrast, we did observe the expected increase in editing frequency ^31^over the course of erythroid differentiation as measured by digital PCR (dPCR) in the Cas9/AAV condition (**Figure S5D**; *P* ≤ 0.0036). However, total cell counts in Cas9/AAV-edited conditions were even lower than control-edited conditions, a finding that was more pronounced at the higher AAV MOI (**Figure 3E-F** and **S5E**; *P* < 0.0001). These data suggest that the Cas9/AAV editing platform may impair HSPC fitness, consistent with prior reports that Cas9-mediated DSBs and AAV6 donor delivery can negatively affect HSPC viability, stemness, and engraftment potential^34,36^.

Finally, to assess whether multiplex editing was able to enhance reactivation of γ-globin and increase output of HbF-producing erythroid cells, we performed hemoglobin tetramer HPLC at Day 14 of differentiation. Because CD34^+^ HSPCs were isolated from healthy adult donors, adult hemoglobin (HbA) was the predominant tetramer while HbF comprised only ∼10% of all hemoglobins in our CBE control-edited condition (**Figure S6A**). Consistent with original Cas9 editing at *BCL11A* and recent clinical reports of base editing at *HBG* (**Table S1**)^12^, single CBE edits at *HBG*, *BCL11A*, or Casgevy increased HbF 3–5-fold from baseline. In contrast, dual *HBG*+*BCL11A* editing produced the strongest induction, exceeding a 5-fold increase (**Fig. 3G**; *P* < 0.0001). Notably, multiplexing corrective edits with *tEPOR* further enhanced HbF production, increasing HbF reactivation by approximately 1.4-1.8-fold relative to matched conditions without *tEPOR* (**Figure 3H-J** and **S6A-C**; *P* ≤ 0.025). In fact, HbF levels reached a mean of 67.2% of total hemoglobin in our triple-edited condition—exceeding prior reports (**Figure S6A** and **Table S1**)—alongside a greater than four-fold increase in cell count (**Figure 3I**). Finally, to mimic post-transplant competition, we performed spike-in experiments and found that all base editing conditions (*BCL11A* CBE, *HBG* CBE, and dual *HBG+BCL11A* CBE) showed even greater increases in HbF when combined with *tEPOR*, with % HbF rising by more than 2-fold relative to the corresponding -*tEPOR* groups (**Figure 3H** and **S6B**; *P* < 0.0001). In contrast, combining Casgevy with *tEPOR* produced a modest, non-significant increase in % HbF relative to Casgevy alone, consistent with inefficient co-occurrence of edits and/or the burden of multiple DSBs.

Together, these results show that integrating *tEPOR* with therapeutic base edits maintains high viability, preserves differentiation, sustains on-target editing efficiency, and substantially enhances both the expansion and functional output of edited erythroid cells.

### Multiplex editing preserves in vitro multi-potential of HSPCs

To assess the clonogenic capacity and lineage potential of edited HSPCs, we performed colony-forming unit (CFU) assays following base editing. Overall colony output was enhanced in base-edited conditions, with *HBG*-edited and multiplex-edited (*HBG+BCL11A+tEPOR*) HSPCs yielding increased colony numbers relative to unedited Mock controls (**Figure S7A**). Notably, Casgevy-edited cells also produced robust colony numbers, whereas Cas9/AAV6-edited cells (at high AAV MOI) showed markedly reduced colony formation. Consistent with these observations, total cell output derived from individual colonies was substantially higher in multiplex-edited HSPCs compared with Mock controls (**Figure S7B**), suggesting enhanced proliferative capacity. Importantly, analysis of colony subtype composition revealed preservation of multilineage differentiation potential across all edited conditions (**Figure S7C**). We also observed a notable increase in the number of BFU-E colonies among the *tEPOR* multiplex-edited condition, while producing comparable numbers of other colonies relative to unedited control HSPCs, consistent with prior studies^21^.

To determine whether corrective edits co-occur with the *tEPOR* edit within the same cells following multiplex editing, we genotyped individual CFU colonies. Across colonies derived from multiplex-edited HSPCs, we detected robust C-to-T editing at the *HBG*, *BCL11A*, and *tEPOR* loci at levels comparable to single edit controls, with sequencing confirming heterozygous or homozygous edits at all three sites across all clones (**Figure S7D**-**F**). Together, these results indicate that multiplex base editing preserves HSPC function while enabling efficient, coordinated editing.

### Multiplex base editing enhances HbF induction in SCD HSPCs

We next assessed whether combinatorial base-editing could enhance erythroid output and HbF induction in the context of SCD. To test this, CD34⁺ HSPCs derived from SCD patients were edited using single-, dual-, or triple-base editing targeting *HBG*, *BCL11A*, and *tEPOR*, with Casgevy included as a clinical benchmark. As in healthy donor HSPCs, post-editing viability and erythroid differentiation efficiency remained high across all conditions (**Figure 4A-B**), indicating that multiplex base editing is well tolerated and does not impair the survival or maturation capacity of SCD HSPCs. Quantification of editing frequencies at Day 4 of differentiation demonstrated consistently high editing across all loci, with multiplexing of two or three CBEs resulting in only minimal reductions compared to single-target editing at the same sites (**Figure 4C** and **S8A**).

**Fig. 4:**
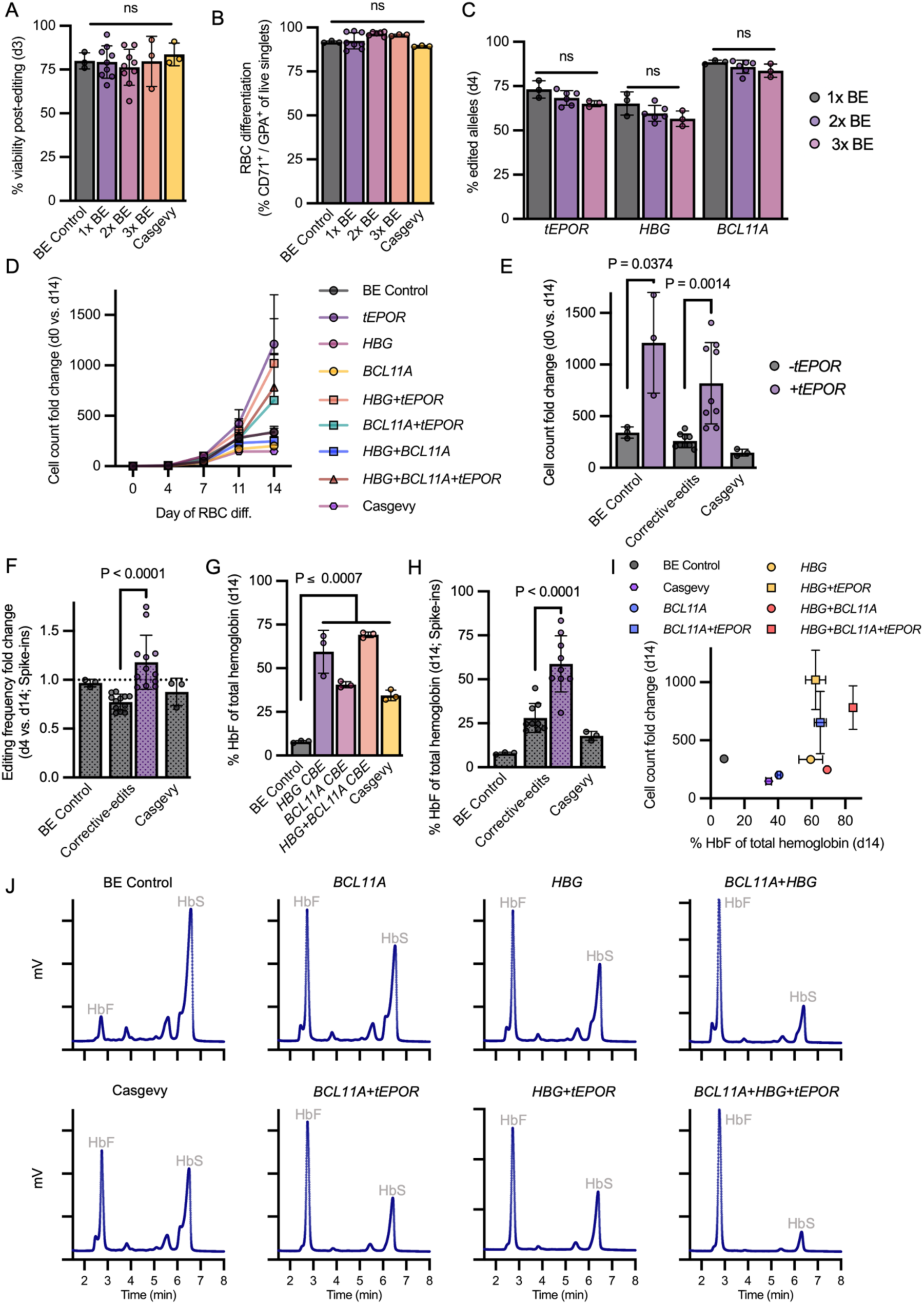
Combinatorial base editing enhances erythroid proliferation and HbF induction in SCD HSPCs. **A.** Percentage of viable cells at Day 3 post-editing across single-, dual-, and triple-BE conditions, Casgevy (*BCL11A* Cas9), and BE control (*CCR5* CBE edit). **B.** Erythroid differentiation efficiency of edited HSPCs at end of erythroid differentiation. **C.** Editing frequencies at Day 4 of erythroid differentiation at the *tEPOR, HBG,* and *BCL11A* loci across single-, dual-, and triple-editing conditions. **D.** Cell count fold change dynamics over course of erythroid differentiation. **E.** Total cell count fold change over course of erythroid differentiation across control (*CCR5* CBE), corrective base-edit (*HBG*, *BCL11A*, or *HBG* + *BCL11A*), and Casgevy conditions, with or without *tEPOR* multiplexing. **F.** Editing frequency fold change over course of erythroid differentiation with spike-in of unedited cells, normalized to control. **G-H.** Percentage of HbF among total hemoglobin following erythroid differentiation, determined by hemoglobin tetramer HPLC. Shown comparing control vs. all single-edited conditions, no spike-in (H) and with spike-in (H). **I.** Relationship between HbF induction and erythroid expansion across editing conditions. Error bars denote ± SEM. **J.** Representative hemoglobin tetramer HPLC chromatograms from cell lysates; fetal hemoglobin (HbF) and sickle hemoglobin (HbS) peaks labeled. Gray columns depict samples without *tEPOR*, and purple columns depict samples with *tEPOR* co-editing. Columns depict mean ± SD. Columns with dotted shading indicate data from spike-in experiments. Statistical significance was assessed using one-way ANOVA with Dunnett’s multiple comparisons test (A-C,G) or unpaired t-tests with Holm-Šidák correction (E,F,H). ns, not significant (*P* > 0.05). Data are shown as replicate measurements from a single donor (n = 1), with three replicates per condition.

We next evaluated the effect of *tEPOR* co-editing on erythroid expansion. When introduced alongside single edits at *HBG* or *BCL11A*, or dual *HBG*+*BCL11A* editing, *tEPOR* increased erythroid expansion relative to both BE controls and matched -*tEPOR* samples, with the proliferative advantage emerging early and increasing over time (**Figure 4D-E** and **S8B**; *P* ≤ 0.037). Cultures harboring corrective CBE edits (*HBG*, *BCL11A*, or their combination) together with *tEPOR* expanded substantially more than Casgevy-edited cells despite comparable initial editing efficiencies (**Figure 4D**-**E** and **S8A**-**B**). Consistent with observations in healthy donor HSPCs and suggestive of ineffective erythropoiesis, both Cas9 and CBE *BCL11A*-edited conditions yielded fewer erythroid cells relative to all other treatments and controls (**Figure S8C**; *P* ≤ 0.0066). Reinforcing the observed cell count increases among all *tEPOR*-edited conditions, we observed increased edited cell frequencies over the course of erythroid differentiation across all *tEPOR* dual- and triple-editing conditions (**Figure 4F** and **S8D**; *P* ≤ 0.029).

Finally, we evaluated whether the enhanced expansion conferred by *tEPOR* improved therapeutic HbF reactivation in the context of SCD. We found that Casgevy increased % HbF from a mean of 7.9% in control-edited cells to 34.5%, while single CBE edits at *HBG* or *BCL11A* exceeded these levels, yielding 59.5% and 40.6% HbF, respectively, and dual *HBG*+*BCL11A* edits increased levels further to 69.2% (**Figure 4G**; *P* ≤ 0.0008). Notably, we found that multiplexing with *tEPOR* further elevated HbF levels (*P* < 0.0001) across all CBE corrective-editing conditions (**Figure 4H-J** and **S8E**), with *HBG*+*BCL11A*+*tEPOR* achieving 84.6% HbF. These levels substantially exceed the ∼60% HbF reported for the most advanced clinical base editing trial (BEAM-101)^37^, while also yielding a >3-fold increase in total cell output (**Figure 4I**). In spike-in experiments designed to mimic post-transplant competition, all CBE-based corrective edits yielded even more pronounced increases in HbF when combined with *tEPOR* (**Figure 4H** and **S8F**; *P* < 0.0001).

These findings support a cooperative interaction between proliferative and corrective edits that simultaneously improves erythroid output and therapeutic hemoglobin production in the context of SCD.

### Multiplex base editing drives expansion of HbF-expressing erythroid cells in β-thalassemia

As in SCD, we next tested whether multiplex base editing could similarly improve erythroid output in β-thalassemia major (BTM) HSPCs. HbF reactivation represents a clinically validated therapeutic approach for both disorders, with Casgevy recently receiving FDA approval for β-thalassemia^38^ following initial approval for SCD. As observed with healthy donor and SCD patient-derived HSPCs, dual- and triple-base editing across *HBG*, *BCL11A*, and *tEPOR* target loci did not adversely affect cell fitness, post-editing viability, or erythroid differentiation efficiency relative to BE control and Casgevy conditions (**Figure 5A-B**). We again observed efficient multiplex editing, albeit with a reduction in editing frequency at *tEPOR* and *HBG* in dual- and triple-editing conditions relative to single-locus editing (**Figure 5C** and **S9A**). Over the course of erythroid differentiation, addition of the *tEPOR* edit enhanced erythroid expansion, with *tEPOR*-containing cultures expanding more than their -*tEPOR* counterparts across all editing conditions, including Casgevy (**Figure 5D**-**E**; *P* ≤ 0.0004). This increased expansion among +*tEPOR* conditions ultimately yielded a greater number of erythroid cells harboring corrective edits (**Figure 5F**). Hemoglobin tetramer HPLC revealed the highest HbF reactivation in *HBG+BCL11A* conditions, both with and without *tEPOR*, while inclusion of *tEPOR* increased the total number of HbF-reactivated erythroid cells (**Figure 5G** and **S9B**-**D**). Further HPLC analysis of individual globin chains showed an increase in γ-globin production among the triple-base editing condition relative to BE control or Casgevy-edited cells (**Figure 5H** and **S9E**-**F**). Together, these results demonstrate that multiplex base editing of *HBG* and *BCL11A* efficiently reactivates HbF in BTM HSPCs, with *tEPOR* co-editing enhancing erythroid expansion without compromising viability or differentiation. This combined strategy ultimately outperforms Casgevy in both erythroid yield and HbF reactivation.

**Fig. 5:**
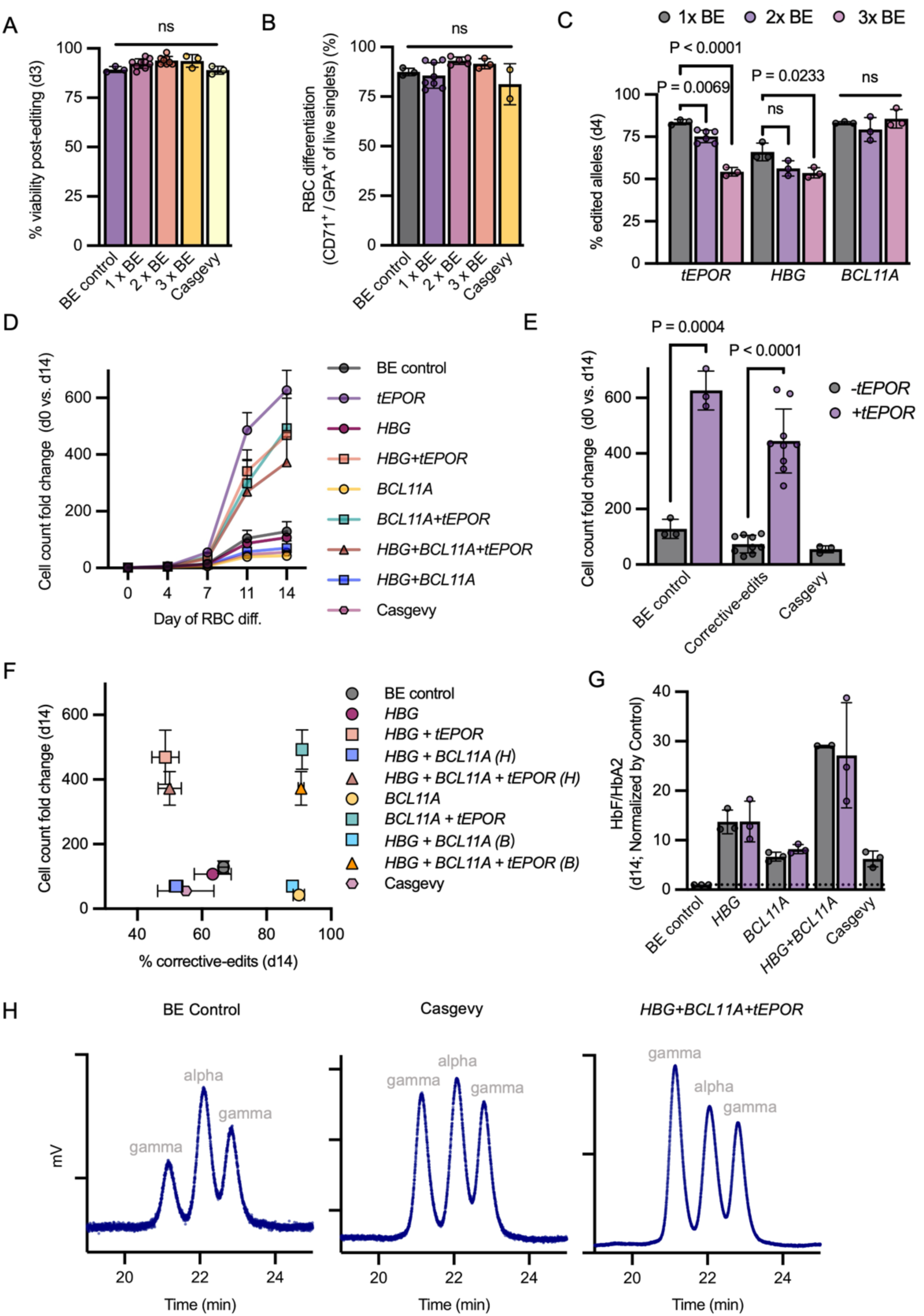
Combinatorial base editing enhances erythroid proliferation and HbF induction in BTM HSPCs. **A.** Percentage of viable cells at Day 3 post-editing across single-, dual-, and triple-BE conditions, Casgevy (*BCL11A* Cas9), and BE control (*CCR5* CBE edit). **B.** Erythroid differentiation efficiency of edited HSPCs at end of erythroid differentiation. **C.** Editing frequencies at Day 4 of erythroid differentiation at the *tEPOR, HBG,* and *BCL11A* loci across single-, dual-, and triple-editing conditions. **D.** Cell expansion dynamics over course of erythroid differentiation. **E.** Total cell count fold change over course of erythroid differentiation across control (*CCR5* CBE), corrective base-edit (*HBG*, *BCL11A*, or *HBG* + *BCL11A*), and Casgevy conditions, with or without *tEPOR* multiplexing. **F.** Correlation between editing frequency and erythroid expansion at end of erythroid differentiation. showing % correct edits (*HBG or BCL11A*) versus cell count fold change. Points are plotted as mean values, with horizontal and vertical error bars indicating ± SD for editing efficiency and expansion, respectively. Conditions are color-coded as indicated, measurements for *HBG* (H) and *BCL11A* (B) target sites are shown separately where applicable. Error bars exceeding the plotting range are truncated. **G.** Ratio of HbF/HbA2 ratio at end of erythroid differentiation, quantified by hemoglobin tetramer HPLC and normalized to control. **H.** Globin chain HPLC chromatograms from cell lysates; gamma- and alpha-globin peaks labeled. Gray columns depict samples without *tEPOR*, and purple columns depict samples with *tEPOR* co-editing. Columns represent mean ± SD. Statistical significance was assessed using one-way ANOVA with Dunnett’s multiple comparisons test (A-C) or unpaired t-tests with Holm-Šidák correction (E). ns, not significant (*P* > 0.05). Data are shown as replicate measurements from a single donor (n = 1), with three replicates per condition.

### Multiplex base-edited HSPCs retain long-term engraftment and multilineage reconstitution in vivo

To assess the impact of combinatorial base editing on long-term engraftment and multilineage reconstitution of HSPCs, we performed xenotransplantation experiments and benchmarked against unedited, Casgevy-, and Cas9/AAV-edited HSPCs (at both medium and low AAV doses). To do so, we edited human CD34⁺ HSPCs with *HBG* (1x), *HBG*+*tEPOR* (2x), or *HBG*+*BCL11A*+*tEPOR* (3x) using a CBE and transplanted cells into NBSGW mice^39^ and analyzed bone marrow composition 16 weeks post-transplantation (**Figure 6A** and **Table S3**). Mice receiving HSPCs edited with medium Cas9/AAV dose (2,000 MOI of in-house AAV without AZD7648 integration enhancer) exhibited low engraftment, with a mean of 1.2% human hematopoietic cell chimerism^40^. This was markedly improved to 49.1% chimerism by reducing the AAV dose to 625 MOI, using commercially purified AAV, and adding AZD7648^33^. In comparison, Casgevy-treated mice displayed intermediate engraftment (∼35-50%). On the other hand, multiplex base-editing conditions achieved higher levels of engraftment, with the 1x BE and 2x BE cohorts reaching ∼60-75% chimerism, and the 3x BE cohort maintaining ∼50-60% chimerism (**Figure 6B, S10**, **S11**, and **S12A**; *P* ≤ 0.025). Lineage analysis revealed preservation of both myeloid and lymphoid lineages that was comparable across all treatments (**Figure 6C**-**D**, **S10**, **S12B**-**C**, and **S13**). These findings are consistent with our *in vitro* CFU data and supports a model in which *tEPOR* promotes expansion of erythroid-committed cells without skewing multi-lineage output.

**Fig. 6:**
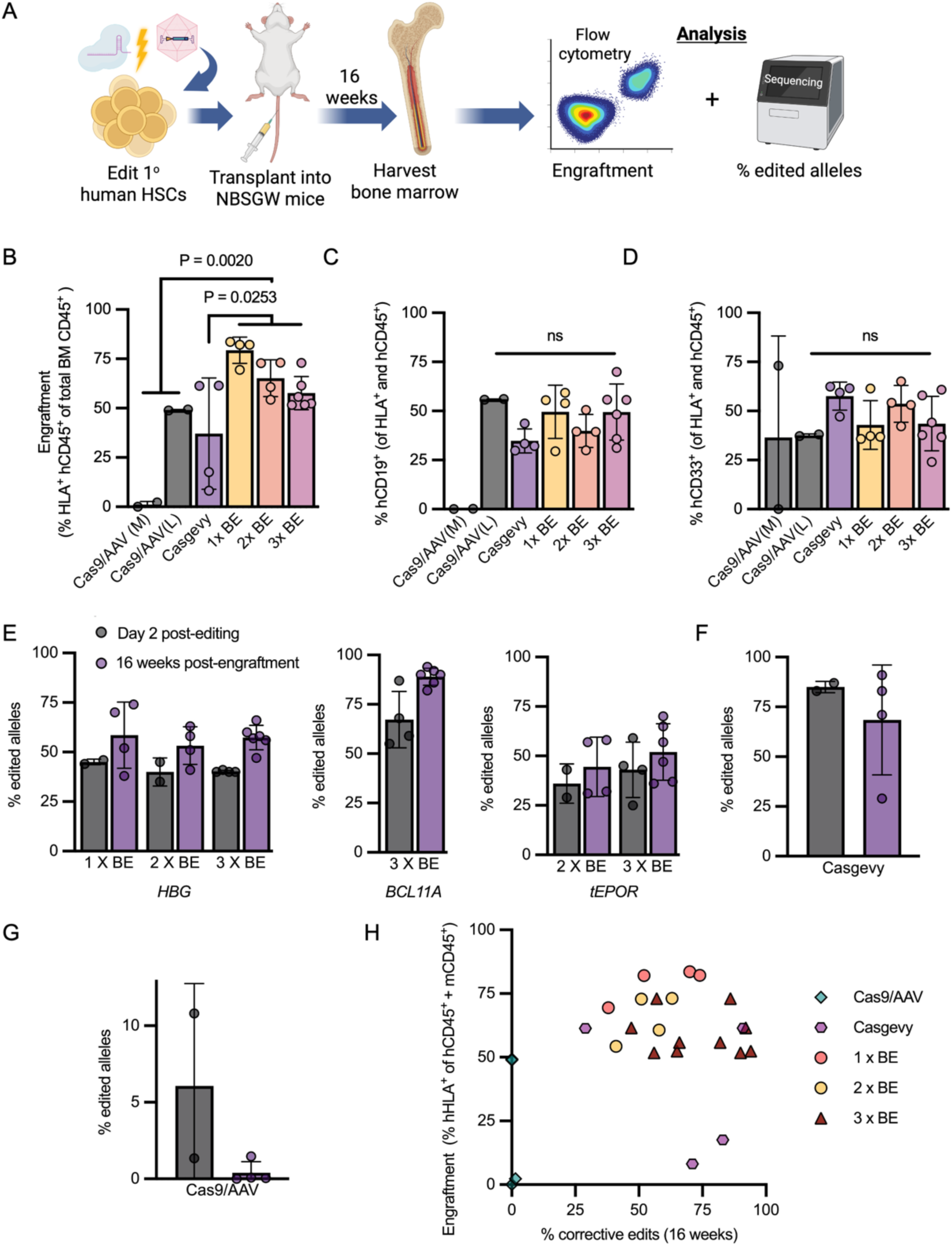
Combinatorial base-edited HSPCs maintain long-term engraftment and multilineage reconstitution. **A.** Schematic of the experimental workflow. Human CD34⁺ HSPCs were edited with single- (*HBG*), dual- (*HBG* + *tEPOR*), or triple- (*BCL11A* + *HBG* + *tEPOR*) BE, as well as Casgevy (*BCL11A* Cas9) or Cas9/AAV6 (integrating *HBB*-IRES-*tEPOR* at *HBA1* locus; Cas9/AAV(L) = MOI 625 + AZD7648; Cas9/AAV(M) = MOI 2,000). HSPCs were then transplanted into sub-lethally irradiated NBSGW mice, and bone marrow was harvested 16 weeks post-transplantation to assess human cell engraftment, lineage distribution, and editing frequencies. **B.** Percentage of human hematopoietic cells in the mouse bone marrow at 16 weeks post-transplantation. **C-D.** Percentage of human lymphoid (C) and myeloid (D) cells among engrafted human hematopoietic population at 16 weeks post-transplantation. **E–G.** Editing frequencies measured 2 days post-editing and 16 weeks post-engraftment across BE (E), Cas9 (F), and Cas9/AAV (G) conditions. **H.** Correlation between human cell engraftment and edited allele frequency for the corrective edit at 16 weeks post-transplantation. Points labeled “Day 2 post-editing” represent edited input samples measured 2 days post-editing before transplantation, whereas other points represent individual mice (n = 22) analyzed 16 weeks post-engraftment. Columns represent mean ± SD. Gray columns depict editing frequencies measured 2 days post-editing, and purple bars indicate frequencies measured 16 weeks post-engraftment (E-H). Statistical significance was assessed using one-way ANOVA with Dunnett’s multiple comparisons test (B-D). ns, not significant (*P* > 0.05).

To determine whether editing was stably maintained in long-term repopulating cells, we quantified editing frequencies at Day 2 post-editing and again in engrafted bone marrow cells isolated 16 weeks after transplantation. Initial editing frequencies were similar across all BE conditions, regardless of whether one, two, or three loci were targeted, indicating that multiplex targeting did not reduce editing efficiency at individual sites (**Figure 6E**). Notably, base editing frequencies were not only maintained among human hematopoietic cells in the bone marrow at Week 16 but increased at all sites relative to those measured in transplanted HSPCs. Among 3x BE conditions, this resulted in a mean of 57.3% edited alleles at *HBG*, 89.0% at *BCL11A*, and 52.0% at *EPOR* among HSPCs at Week 16 post-transplant. While preferential engraftment of more highly edited HSPCs is a potential explanation, we speculate that the increase may instead reflect continued editing activity from BE mRNA beyond the Day 2 timepoint when cells were initially assayed. In contrast, despite Cas9 also being delivered as mRNA, editing frequencies declined from a mean of 85.0% edited alleles at *BCL11A* at Day 2 post-editing to 68.5% among cells harvested at Week 16 (**Figure 6F**). Consistent with our observations, clinical studies of Casgevy also reported declines in edited cell frequencies following transplantation^1^. A similar but more pronounced loss of edited alleles was observed among HSPCs targeted using Cas9/AAV, with mean editing of 6.1% at Day 2 falling to 0.4% among cells harvested at Week 16 (**Figure 6G**). While these results suggest that the editing protocol could likely be optimized to achieve a more highly edited input cell population, the decline in editing frequency among long-term repopulating human cells is consistent with prior studies^41–43^ and likely reflects a negative impact of AAV exposure on HSPC viability or stemness, particularly at the higher AAV dose.

Finally, a Sysmex analyzer was used to assess hematologic parameters in mice at Week 16 post-transplant. Measurements of white blood cells, red blood cells, hemoglobin, platelets, neutrophils, and lymphocytes were broadly comparable across treatment conditions, with values clustering near cohort averages. No evidence of clinically meaningful hematologic toxicity was observed following base editing, Cas9, or Cas9/AAV treatments (**Figure S14**). Consistent with known limitations of xenograft models, human erythroid output was not detectable *in vivo*^44,45^. However, multilineage engraftment, together with robust erythroid amplification *in vitro* and CFU assays, supports a model in which *tEPOR* selectively expands erythroid-committed progenitors without diminishing other lineages.

Together, these findings establish multiplex base editing as an effective strategy for installing and maintaining multiple therapeutic edits in long-term repopulating HSPCs while preserving robust engraftment and multilineage reconstitution *in vivo* (**Figure 6H**). In contrast, nuclease- and AAV-based approaches exhibited reduced engraftment and loss of edited alleles over time, highlighting the advantages of base editing for multiplex HSPC engineering.

## Discussion

Genome editing strategies for hemoglobinopathies have historically focused on correcting or compensating for disease-causing mutations. In this study, we demonstrate that multiplex base editing can extend this paradigm by coupling therapeutic edits with lineage-specific fitness engineering to amplify the output of corrected cells. Throughout our work, we found that multiplex base editing was well tolerated in primary human HSPCs. Dual- and triple-editing conditions maintained high editing frequencies, preserved viability, supported normal erythroid differentiation, and retained long-term multilineage engraftment in xenograft models. Clonal analysis confirmed highly coordinated editing across all three target loci, with individual clones carrying heterozygous or homozygous modifications at each target site. Importantly, multiplex base editing compared favorably with nuclease-based strategies such as Casgevy or Cas9/AAV editing platforms, which exhibited reduced viability, lower erythroid output, and decreased long-term retention of edited alleles. These findings highlight a key advantage of base editing for combinatorial genome engineering: the ability to install precise nucleotide substitutions with minimal double-strand DNA breaks, thereby minimizing genotoxic stress while enabling cooperative multi-locus editing.

Leveraging this capability, we combined HbF-reactivating edits with introduction of a naturally occurring *tEPOR* to engineer erythroid lineage amplification. This strategy confers a selective proliferative advantage to edited erythroid progenitors, resulting in both increased erythroid output and progressive enrichment of therapeutically edited cells during differentiation. Notably, *tEPOR* editing also appeared to rescue the reduced erythroid output associated with disruption of the *BCL11A* erythroid enhancer, suggesting that lineage-amplifying edits may mitigate ineffective erythropoiesis that can arise from perturbation of erythroid regulatory networks^46^.

An important feature of multiplex base editing as a platform is the ability to precisely recreate beneficial human genetic variants that have already been characterized in rare individuals. HbF-reactivating mutations associated with hereditary persistence of fetal hemoglobin (HPFH), together with the *EPOR* truncation studied here, represent naturally occurring alleles linked to benign or advantageous hematologic phenotypes. Base editing provides a direct means to install such variants in therapeutic cells while preserving endogenous gene regulation. Combining these naturally occurring alleles through multiplex editing creates opportunities to engineer cooperative genetic programs that do not co-occur in nature but are built from variants with known biological consequences and favorable safety profiles.

These findings have important implications for the clinical translation of genome editing therapies for hemoglobinopathies. Current *ex vivo* editing strategies rely on achieving high levels of engraftment following myeloablative conditioning to reach therapeutic thresholds^2^. However, such regimens remain a major barrier due to toxicity and long-term complications. Antibody-based conditioning strategies that selectively deplete hematopoietic stem cells—such as anti-CD117 monoclonal antibodies^47,48^—have emerged as promising alternatives and are now being evaluated in multiple clinical trials. While these approaches substantially reduce toxicity, they consistently yield lower levels of donor HSPC engraftment compared with conventional chemotherapy-based conditioning. In this context, lineage amplification could compensate for limited HSPC isolation from SCD patients as well as reduced stem cell engraftment achieved with targeted conditioning approaches, potentially enabling therapeutic benefit at lower levels of edited HSPC chimerism.

Beyond the hemoglobinopathies, these results position multiplex base editing as a modular and generalizable platform for combining therapeutic and functional edits within the same cell. In addition to lineage amplification, this approach can be extended to incorporate features such as immune evasion, antigen deletion, or epitope shielding (**Table S2**)^14,49–52^. The ability to layer these modifications within a single editing strategy expands the design space of genome-edited cell therapies and provides a foundation for more sophisticated engineered therapies.

## Acknowledgements

The authors thank the following funding sources that made this work possible: K.J. was supported by the UCSF-California Institute for Regenerative Medicine Scholars Training Program (EDUC4-12812); B.J.L. was supported by the National Science Foundation Graduate Research Fellowship Program; T.C.M. was supported by the UCSF Center for Maternal-Fetal Precision Medicine; M.K.C. was supported by an NIH Director’s New Innovator Award (DP2- HL185102) and UCSF Catalyst Award. We thank Serine Avagyan for assistance in establishing the NGS pipeline and Vinh Nguyen for flow cytometry support at the UCSF Flow Cytometry Core. We acknowledge the Pediatrics/Winship Flow Cytometry Core of the Winship Cancer Institute of Emory University and Children’s Healthcare of Atlanta, supported by NIH/NCI award P30CA138292. We also thank Sofia Luna (Matthew Porteus laboratory at Stanford) for providing the AAV vector encoding the *HBB*-IRES-*tEPOR* cassette targeting the *HBA1* locus.

## Author contributions

M.K.C. supervised the *in vitro* experiments. T.C.M., V.A.S., and M.K.C. supervised the xenotransplantation experiments. M.C.W., and V.A.S. provided the patient-derived cells. K.J., R.S., B.J.L., M.A.P., R.C., V.A.S., and M.K.C. designed the experiments. K.J., E.S., R.S., B.J.L., M.A.P., R.C., E.M.F., Z.K., S.N.C., D.S., X.Y., and M.C. carried out the experiments. K.J., E.S., X.Z., M.A.P., R.C., and M.K.C. analyzed the data. K.J. and M.K.C. wrote the manuscript.

## Supplementary Figures

**Table S1:**
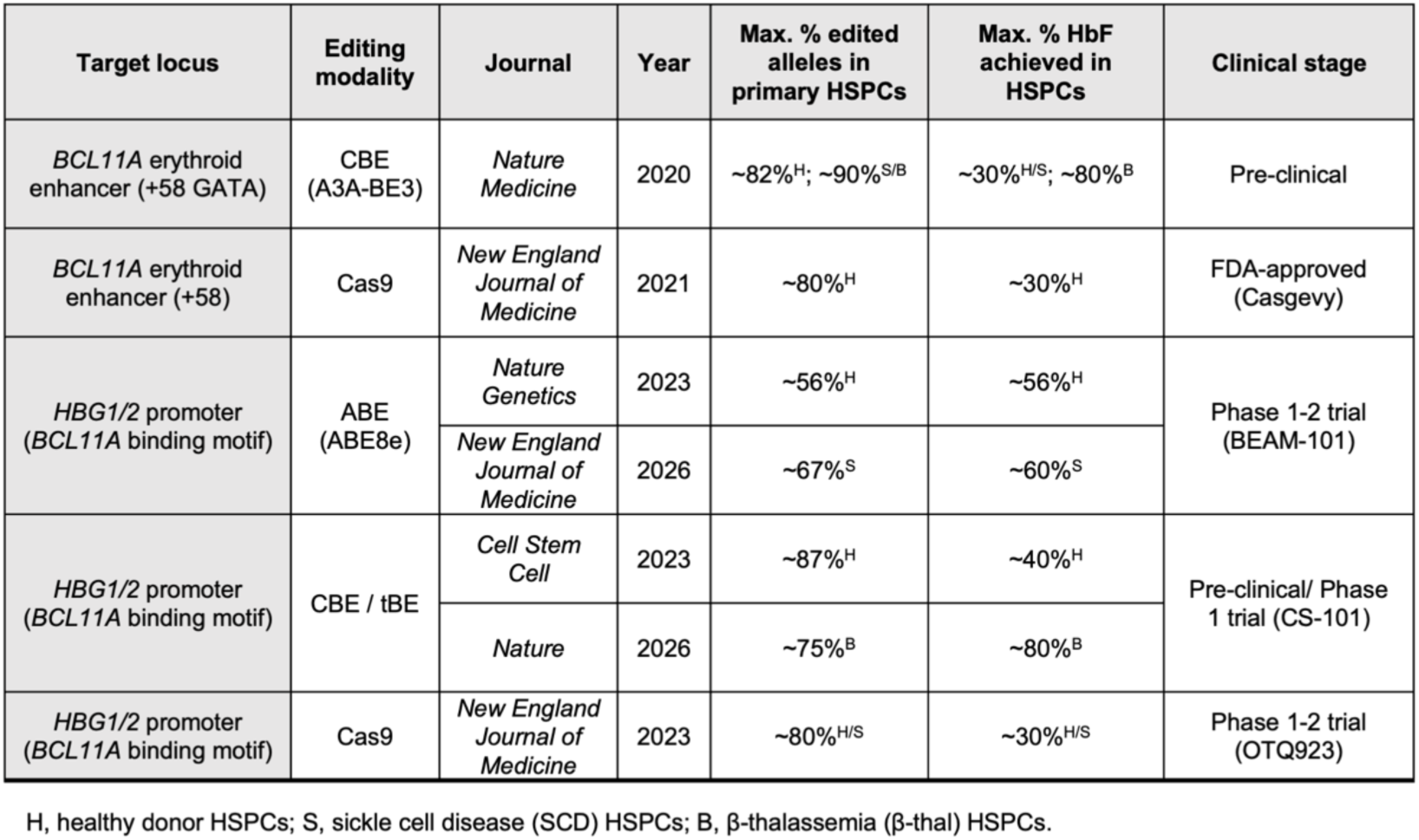
Summary of clinical HbF-reactivating genome editing strategies, related to Figure 3. Table lists target locus, editing modality, journal and year first published, maximum % edited alleles in primary HSPCs, maximum % HbF (of total hemoglobin) achieved, and clinical development stage. tBE: transformer base editor.

**Table S2:**
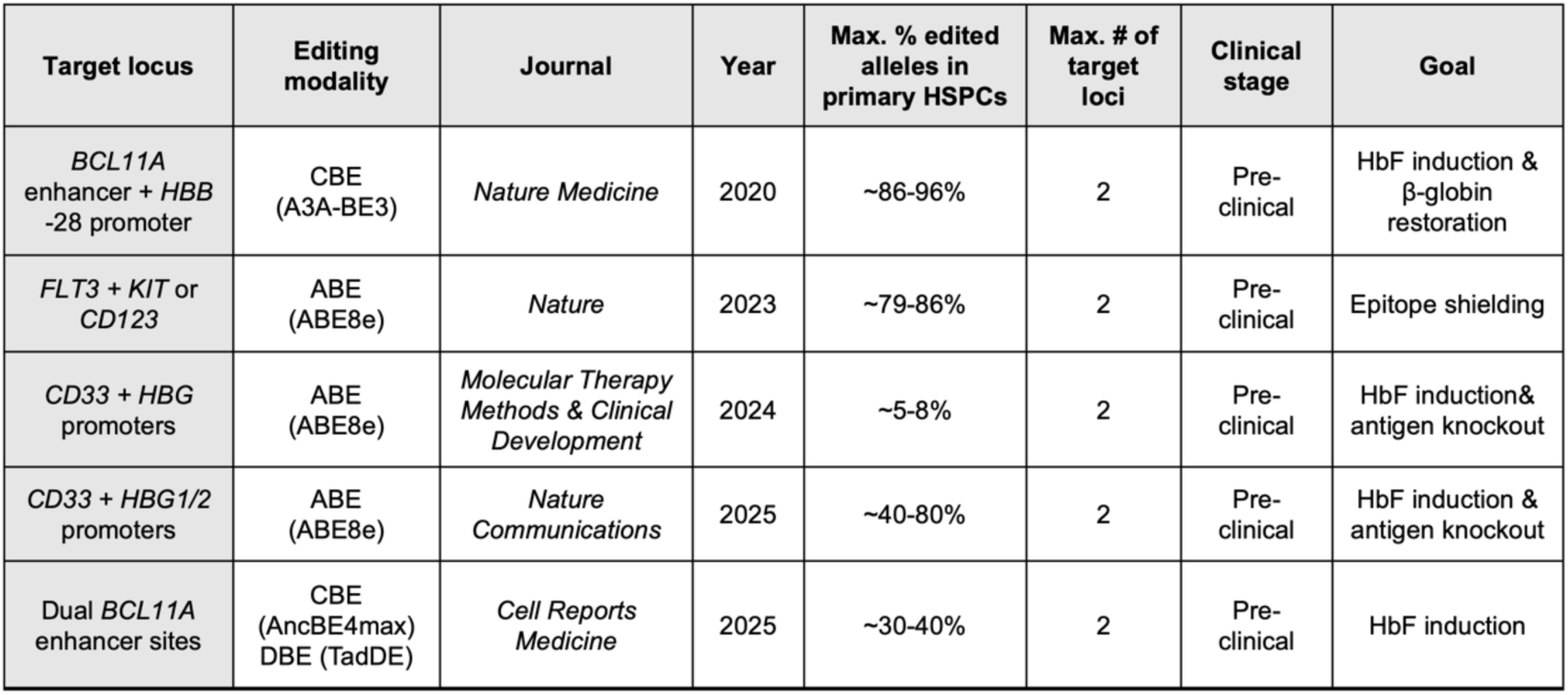
Summary of multiplex base-editing strategies in HSPCs, related to Figure 3. Table lists target locus, editing modality, journal and year published, maximum % edited alleles in primary HSPCs, maximum # of target loci, and clinical development stage.

**Table S3:** Summary of NBSGW mouse transplantation conditions, related to Figure 6. Table lists mouse ID, age at injection, weight at injection, treatment group, and total number of human HSPCs transplanted.

| Mouse ID | Age at injection (wks) | Weight (g) | Treatment group | # HSPCs injected |
| --- | --- | --- | --- | --- |
| 10056 | 8 | 19.9 | Cas9/AAV(M) | 7.50E+05 |
| 10057 | 8 | 18.9 | Cas9/AAV(M) | 7.50E+05 |
| 10058 | 8 | 15.7 | Cas9/AAV(L) | 7.50E+05 |
| 10059 | 8 | 16.8 | Cas9/AAV(L) | 7.50E+05 |
| 10061 | 8 | 19.6 | Casgevy | 7.50E+05 |
| 10062 | 8 | 22.3 | Casgevy | 7.50E+05 |
| 10063 | 8 | 18 | Casgevy | 7.50E+05 |
| 10064 | 8 | 20.3 | Casgevy | 7.50E+05 |
| 10067 | 8 | 21.1 | *HBG + BCL11A + tEPOR* | 7.50E+05 |
| 10068 | 8 | 17.7 | *HBG + BCL11A + tEPOR* | 7.50E+05 |
| 10044 | 12 | 21.9 | *HBG* | 5.00E+05 |
| 10045 | 12 | 24.1 | *HBG* | 5.00E+05 |
| 10046 | 12 | 24.2 | *HBG* | 5.00E+05 |
| 10047 | 12 | 22.0 | *HBG* | 5.00E+05 |
| 10048 | 12 | 22.8 | *HBG + tEPOR* | 5.00E+05 |
| 10049 | 12 | 20.9 | *HBG + tEPOR* | 5.00E+05 |
| 10050 | 12 | 23.0 | *HBG + tEPOR* | 5.00E+05 |
| 10051 | 12 | 21.8 | *HBG + tEPOR* | 5.00E+05 |
| 10052 | 12 | 23.8 | *HBG + tEPOR + BCL11A* | 5.00E+05 |
| 10053 | 12 | 22.6 | *HBG + tEPOR + BCL11A* | 5.00E+05 |
| 10054 | 12 | 22.3 | *HBG + tEPOR + BCL11A* | 5.00E+05 |
| 10055 | 12 | 23.7 | *HBG + tEPOR + BCL11A* | 5.00E+05 |

**Figure S1:**
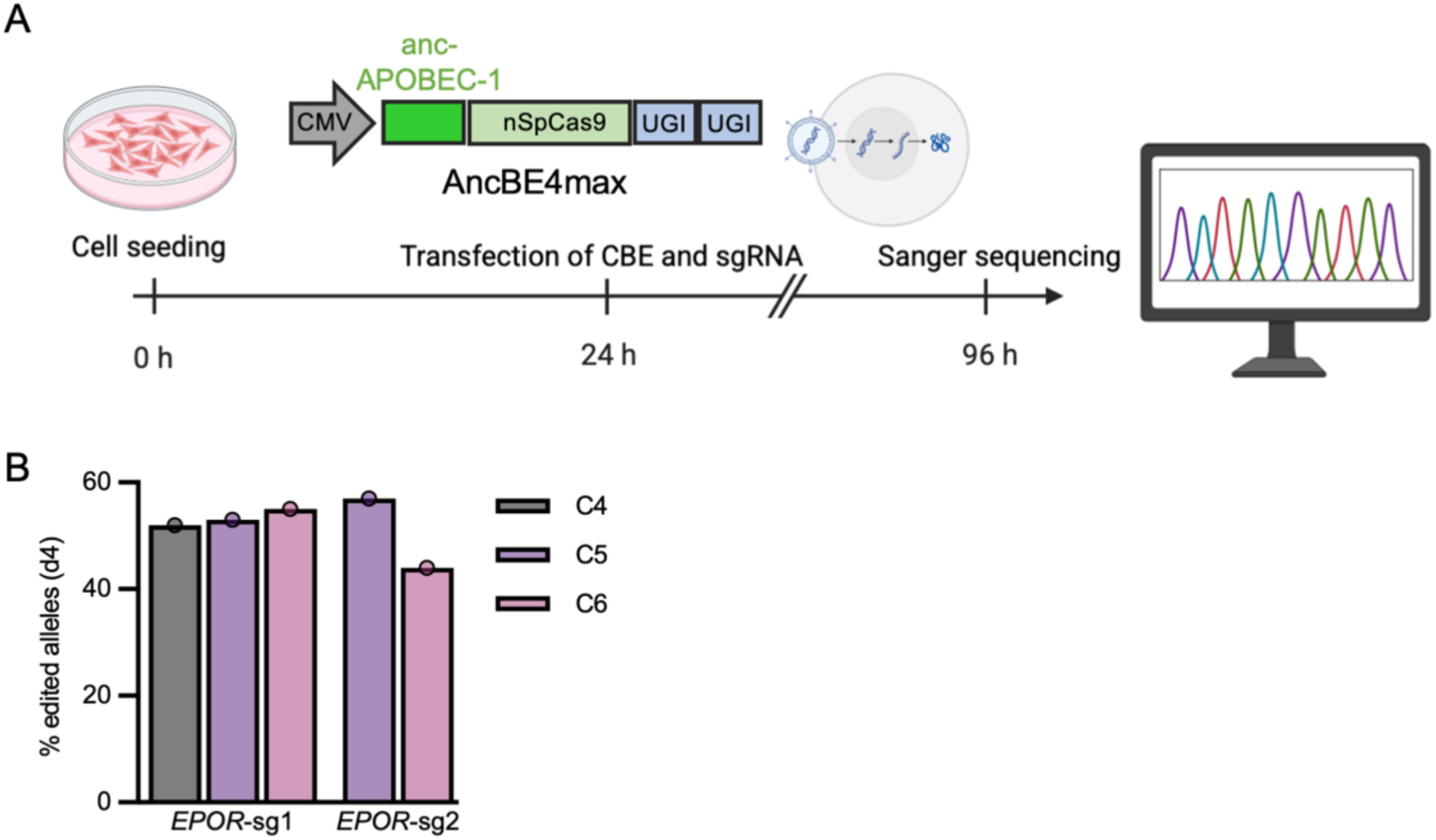
*EPOR* base editing in HeLa cells, related to Figure 1. **A.** Schematic of base editing and analysis workflow in HeLa cells. **B.** Editing frequencies at Day 4 post-transfection determined by EditR analysis. Edited cytosines within the target regions are numbered relative to the nucleotide distal from the PAM.

**Figure S2:**
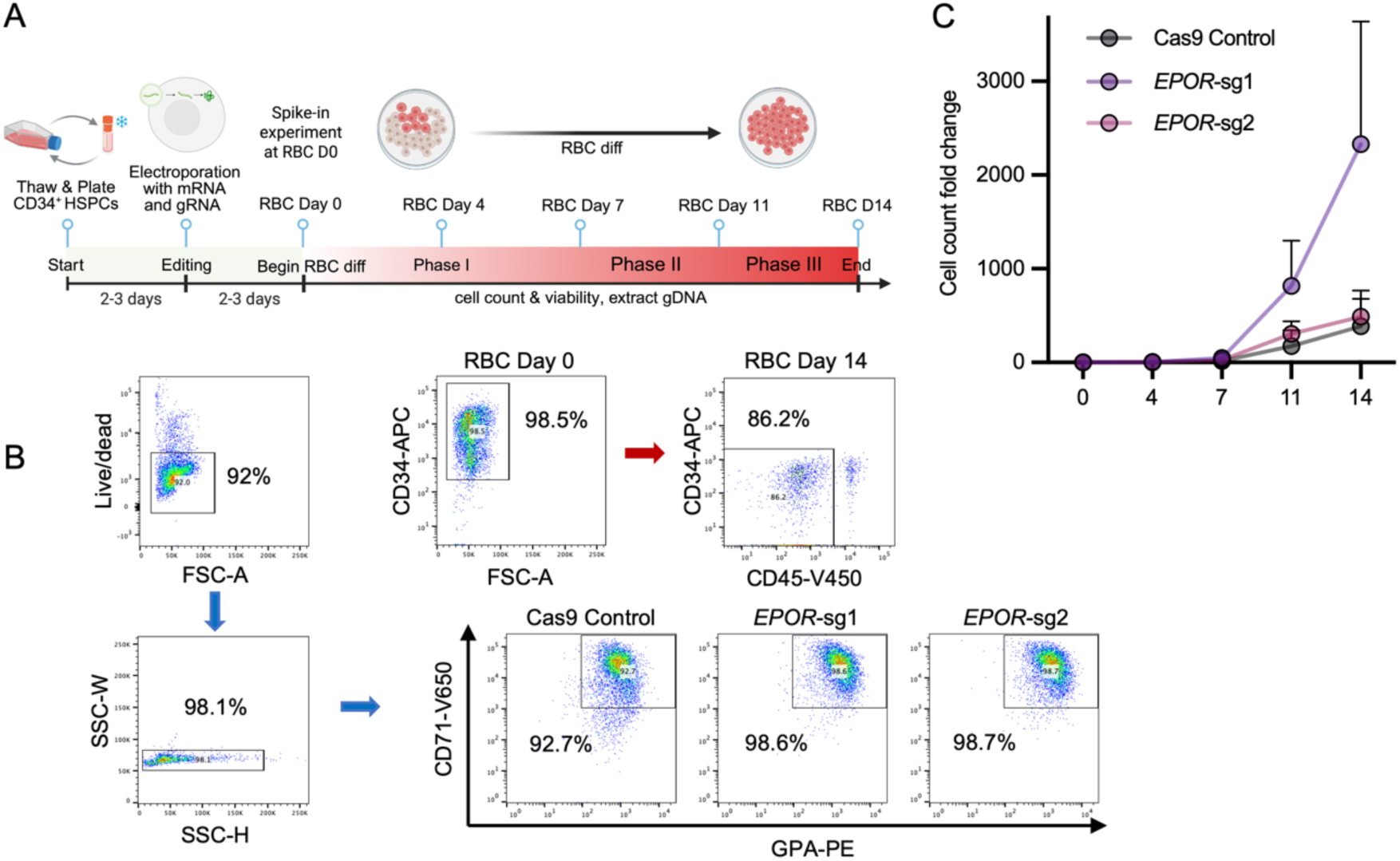
*EPOR* base editing in primary HSPCs, related to Figure 1. **A.** Schematic of base editing workflow and *in vitro* erythroid differentiation of primary human HSPCs. **B.** Representative flow cytometry plots of WT HSPCs at Day 14 of differentiation showing erythroid maturation (CD34^-^/CD45^-^/CD71^+^/GPA^+^). **C.** Cell count fold change of edited HSPCs over the course of erythroid differentiation.

**Figure S3:**
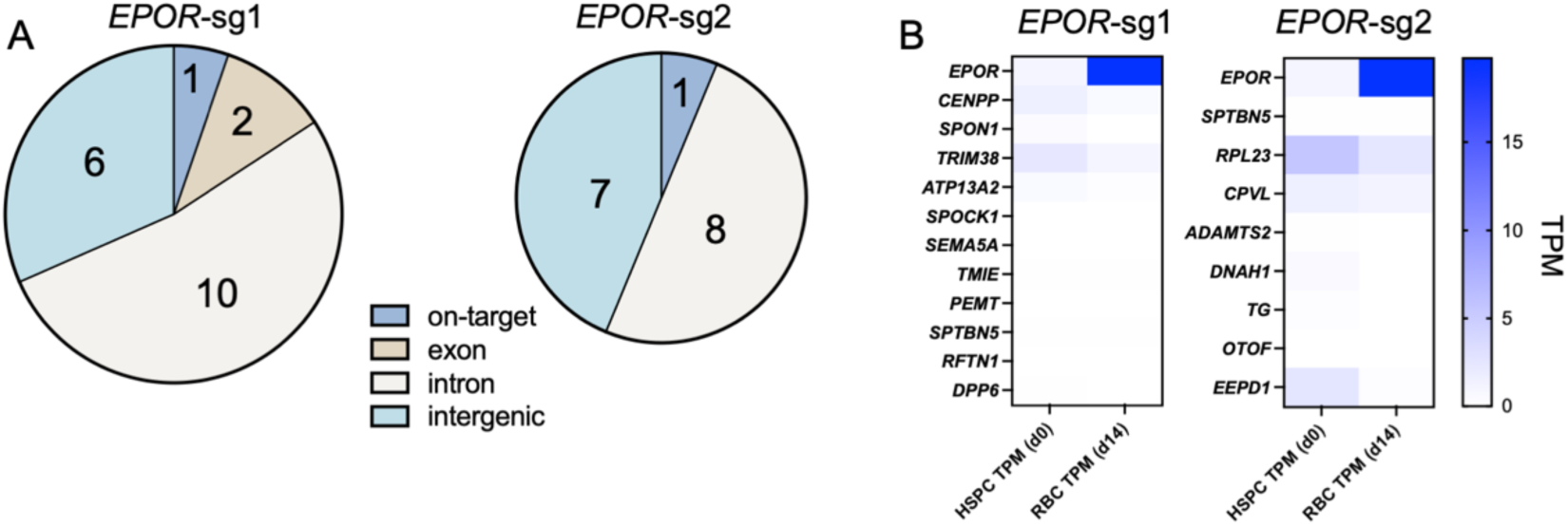
Off-target editing for *EPOR* gRNAs, related to Figure 1. **A.** Distribution of predicted off-target sites across genomic regions for each *EPOR* gRNA. **B.** Expression data (shown as transcripts per million; TPM) for *EPOR* as well as genes with potential off-target editing in exons or introns.

**Figure S4:**
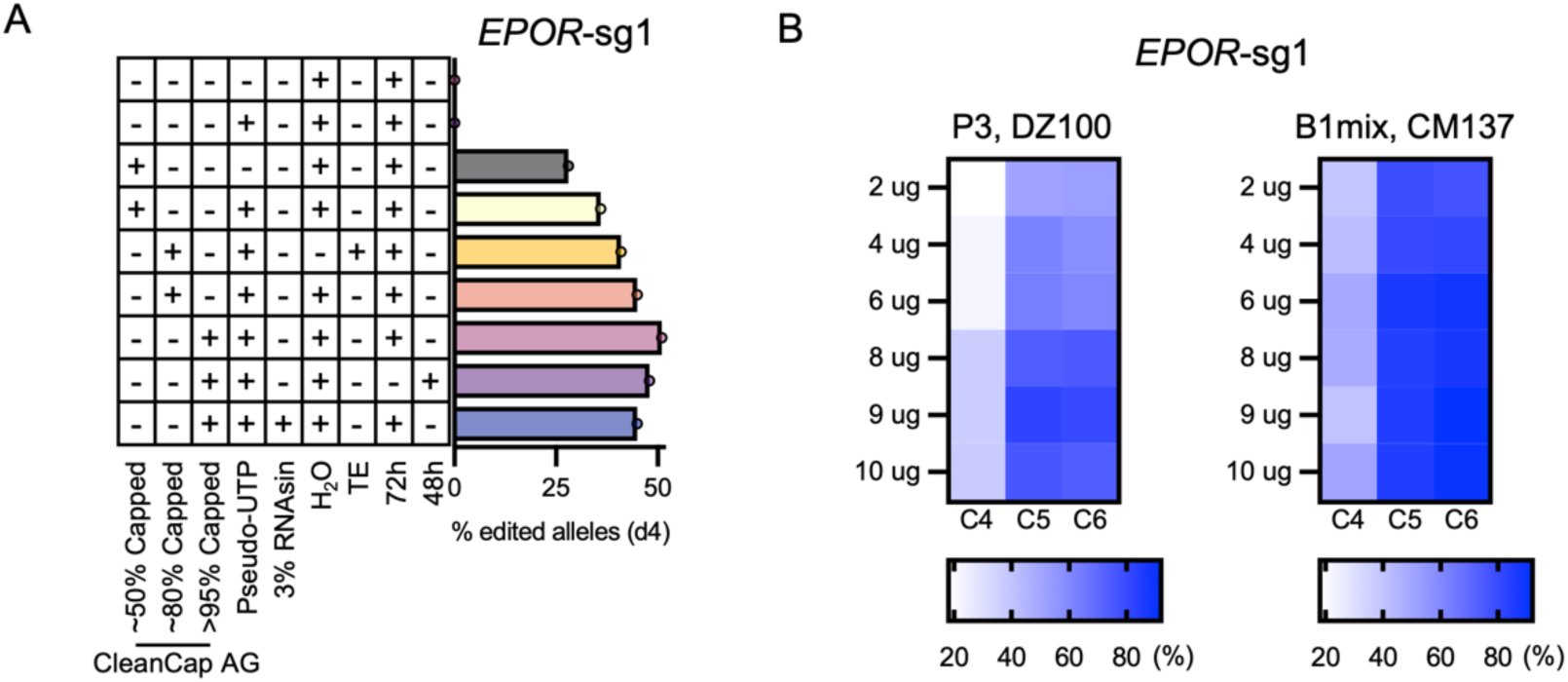
Optimization of base editor mRNA and electroporation in HSPCs, related to Figure 2. **A.** Editing frequencies at the *EPOR-sg1* locus measured at Day 4 post-electroporation with CBE6 mRNA prepared under different conditions, including capping strategies (CleanCap AG), incorporation of modified nucleotides (e.g., pseudouridine), RNase inhibitor addition, freeze-thaw cycles, and storage duration. **B.** Heatmaps showing *EPOR*-sg1 editing frequencies across varying CBE6 mRNA doses (2-10 µg) and electroporation conditions using Lonza 4D nucleofector with P3 buffer + DZ-100 code or B1mix buffer + CM-137 code. Data are shown as replicate measurements from a single donor (n = 1).

**Figure S5:**
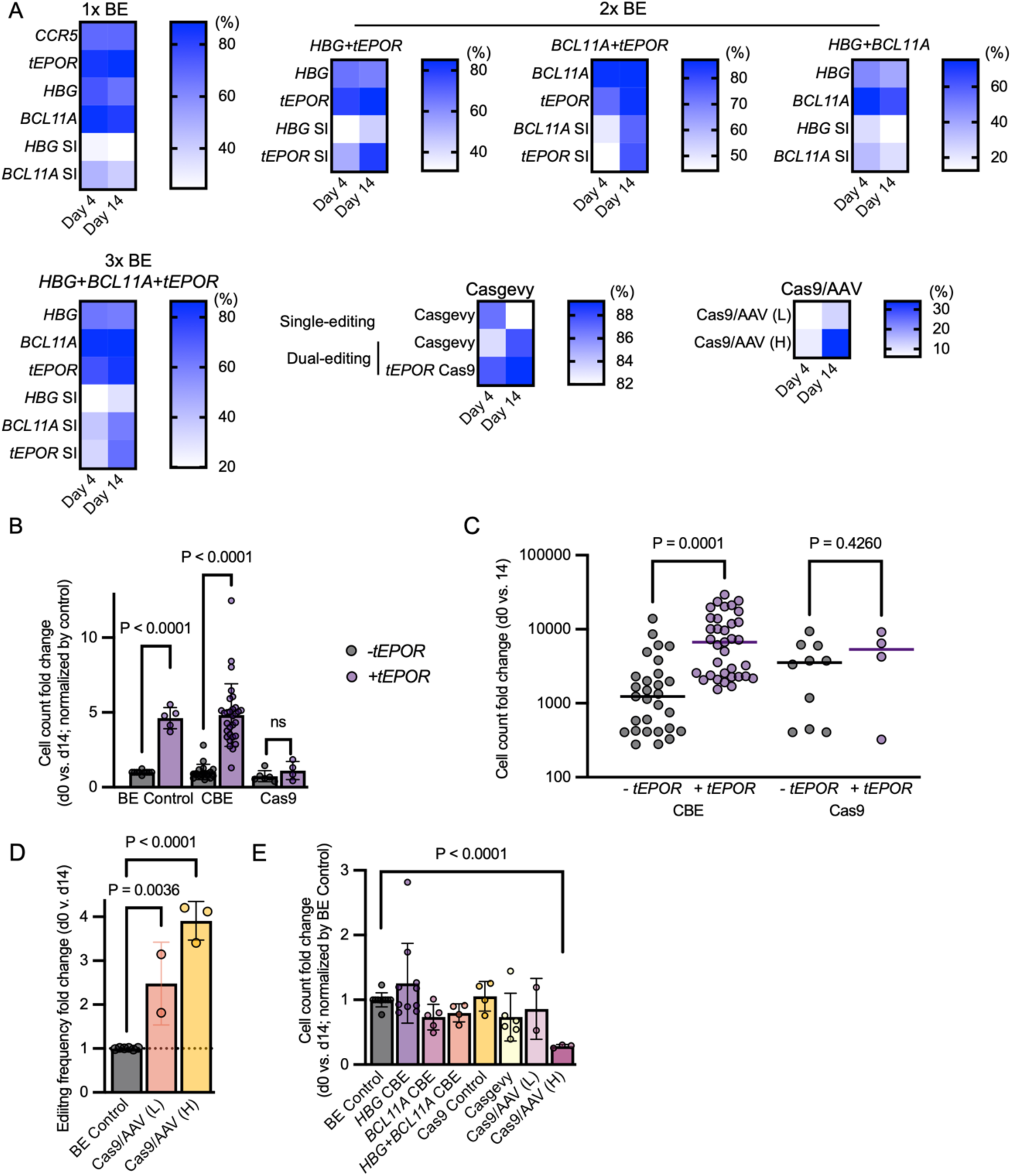
Multiplex base editing combines *tEPOR* with *BCL11A* and *HBG* edits in healthy donor HSPCs, related to Figure 3. **A.** Heatmaps showing editing frequencies at Day 4 and Day 14 of erythroid differentiation. SI indicates spike-in experiments. **B-C.** Total cell count fold change over course of erythroid differentiation among combined base-editing and Cas9-based strategies, normalized to BE control (B) and non-normalized (C). **D.** Editing frequency fold change over course of erythroid differentiation among Cas9/AAV conditions, normalized to control. **E.** Total cell count fold change over course of erythroid differentiation across all conditions, normalized to BE control. All columns depict mean and error bars depict SD. Statistical significance was determined using unpaired t-tests with Holm-Šidák correction (B, C), one-way ANOVA with Dunnett’s multiple comparisons test (D), or linear mixed-effects models (E). ns, not significant (*P* > 0.05). Data are shown as individual measurements from independent donors (n = 7 total), with some donors represented more than once in selected conditions. Sample availability varies across groups, as not all conditions were tested in all donors.

**Figure S6:**
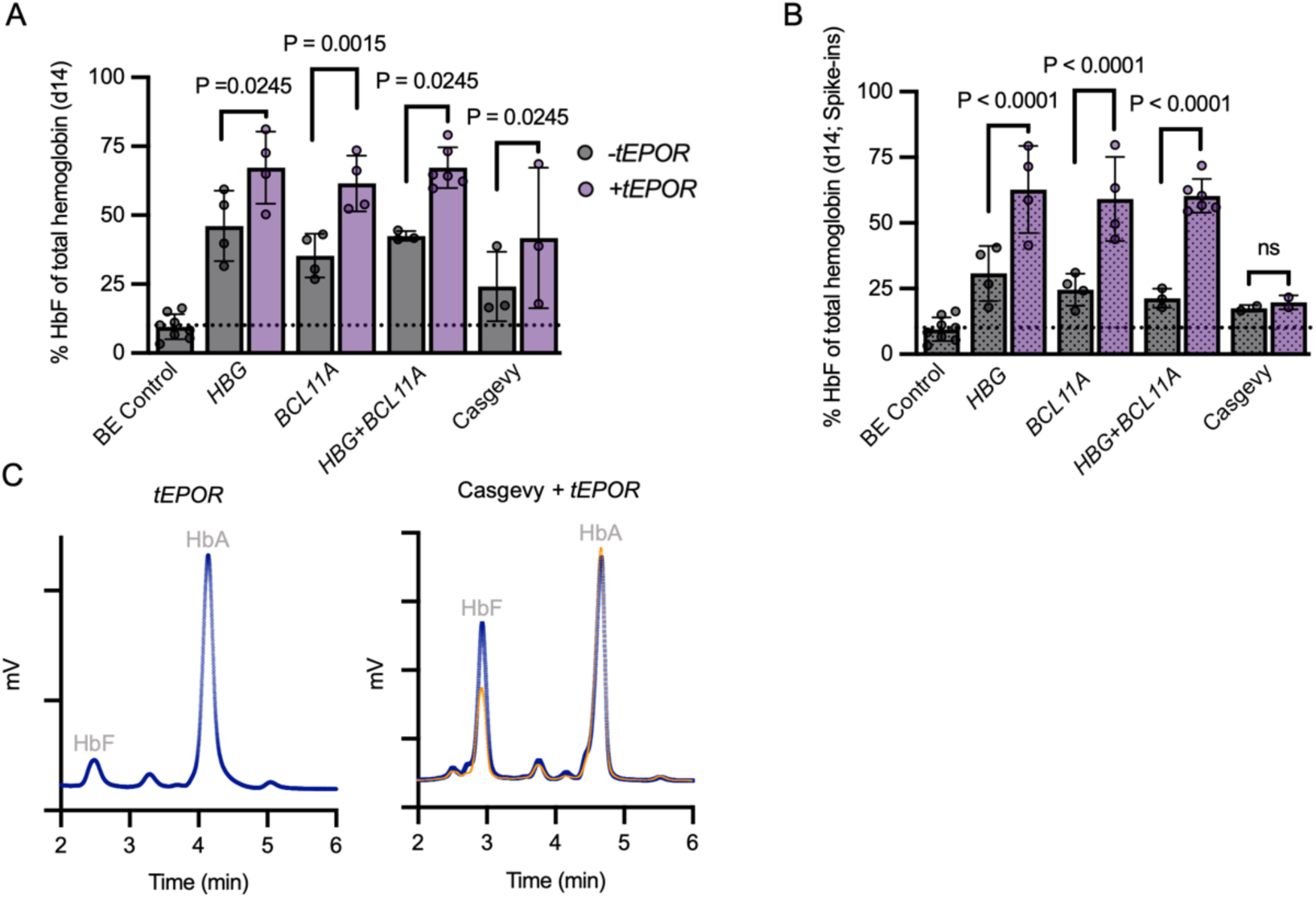
Fetal hemoglobin induction following multiplex base editing in healthy donor HSPCs, related to Figure 3. **A-B.** Percentage of HbF among total hemoglobin following erythroid differentiation, determined by hemoglobin tetramer HPLC, without spike-in (A) and with spike-in (B) of unedited cells. Columns depict mean ± SD. Columns with dotted shading depict spike-in samples. Statistical significance was assessed using a linear mixed-effects model with donor as a random effect and Holm-adjusted multiple comparisons; ns, not significant (*P* > 0.05). Data are shown as individual measurements from independent donors (n = 7 total), with some donors represented more than once in selected conditions. Sample availability varies across groups, as not all conditions were tested in all donors. **C.** Representative hemoglobin tetramer HPLC chromatograms from cell lysates; fetal hemoglobin (HbF) and adult hemoglobin (HbA) peaks labeled. The orange trace depicts Casgevy-alone chromatogram from Fig. 3J overlaid with the Casgevy + *tEPOR* chromatogram for comparison.

**Figure S7:**
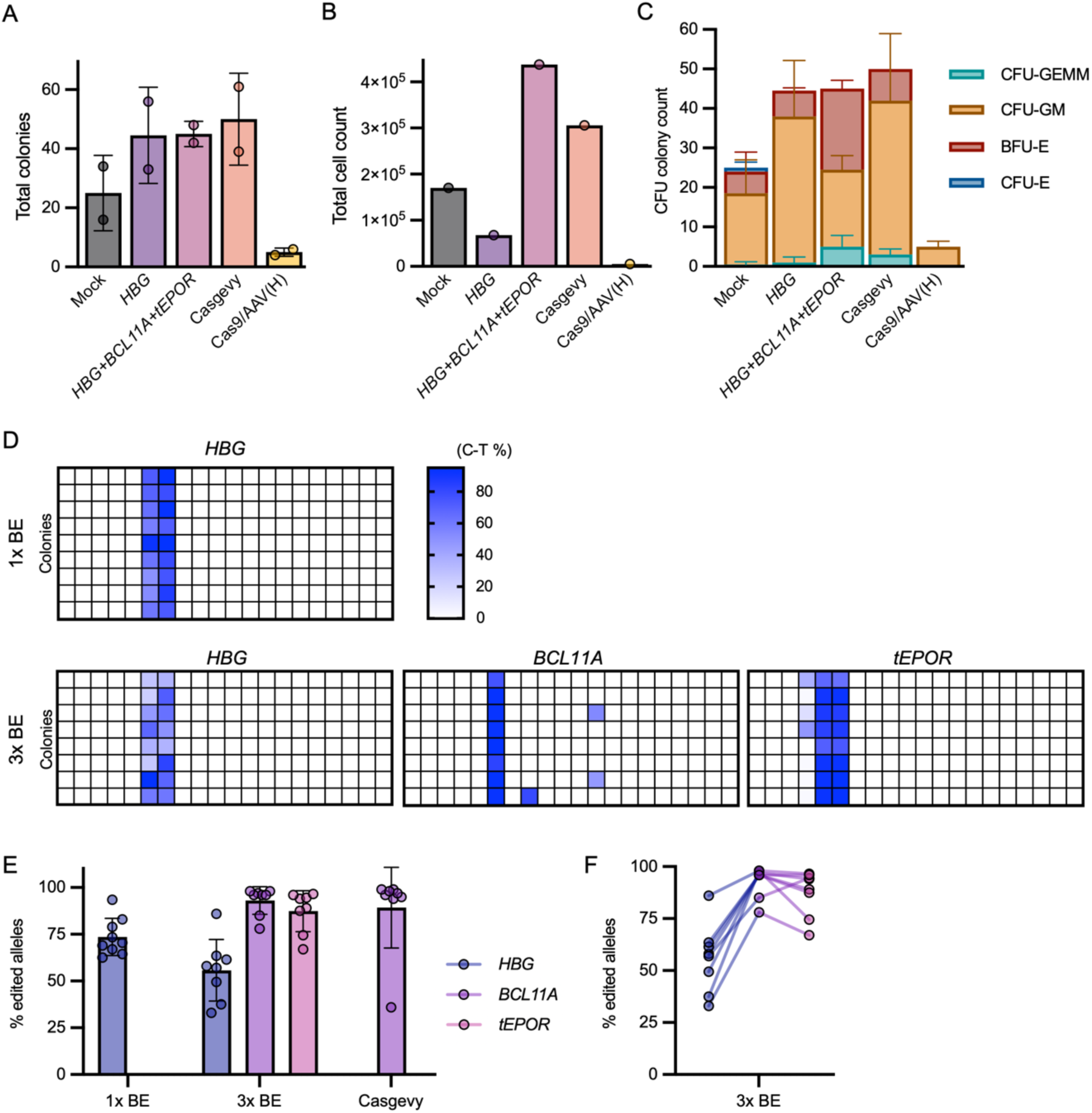
Clonal analysis of base-edited healthy donor HSPCs, related to Figure 3. **A.** Total number of colony-forming units generated across editing conditions. **B.** Total viable cell numbers recovered from each CFU condition. **C.** Distribution of CFU colony subtypes (CFU-GEMM, CFU-GM, BFU-E, and CFU-E). **D.** Genotyping of individual CFU colonies derived from *HBG* single-edited (1x BE) and multiplex-edited (*HBG + BCL11A + tEPOR*; 3x BE) HSPCs. Each row represents one colony, with color indicating C-to-T editing frequency (%) across target loci. **E-F.** Summary of genotyping data generated from individual CFU colonies derived from edited HSPCs across all editing conditions (E) and linked across each target locus within a single colony among 3x BE-edited cells (F). BE conditions were quantified as an average of C-to-T changes at target alleles and Casgevy condition quantifies the percent of alleles with indels. Editing frequencies for base-edited loci were quantified as the average C-to-T conversion across editable cytosines within the base-editing window (and across the highly homologous *HBG1* and *HBG2* loci, which were not distinguished) Columns depict mean ± SD. Data are shown as replicate measurements from a single donor (n = 1).

**Figure S8:**
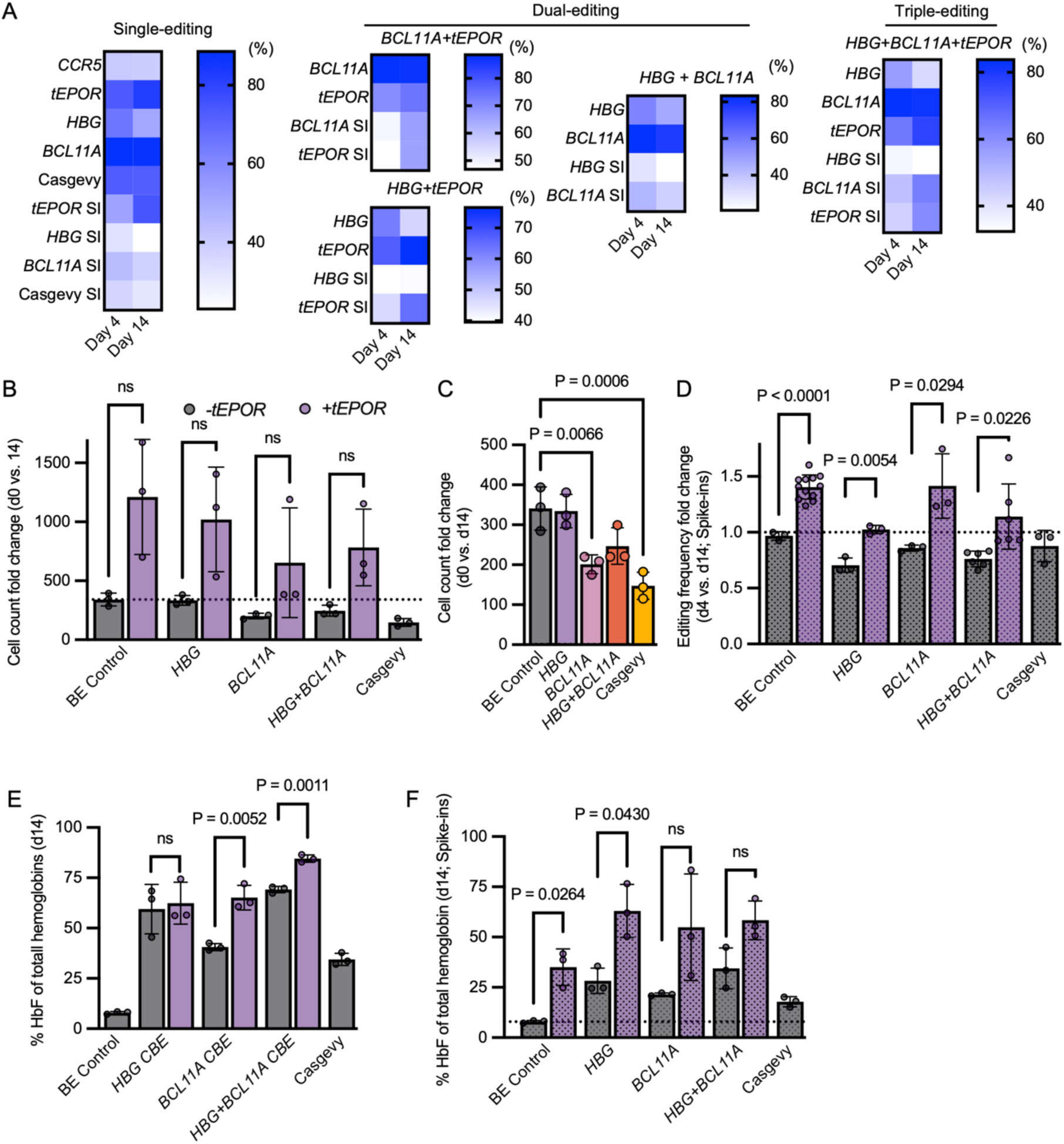
Multiplex base editing in SCD HSPCs, related to Figure 4. **A.** Heatmaps showing editing frequencies at Day 4 and Day 14 of erythroid differentiation. SI indicates spike-in experiments. **B.** Total cell count fold change over course of erythroid differentiation across all conditions. **C.** Total cell count fold change over course of erythroid differentiation across editing conditions compared to BE control. Only statistically significant comparisons are annotated with P values; non-significant comparisons are not shown. **D.** Editing frequency fold change over course of erythroid differentiation with spike-in of unedited cells, normalized to control. Only statistically significant comparisons are annotated with P values; non-significant comparisons are not shown. **E-F.** Percentage of HbF among total hemoglobin following erythroid differentiation, determined by hemoglobin tetramer HPLC, without spike-in (A) and with spike-in (B) of unedited cells. Gray columns depict samples without *tEPOR*, and purple depict samples with *tEPOR* co-editing. Columns depict mean ± SD. Columns with dotted shading depict spike-in samples. Statistical significance was determined using one-way ANOVA with Dunnett’s multiple comparisons test (C) or unpaired t-tests with Holm-Šidák correction (B, D, E-F); ns, not significant (*P* > 0.05). Data are shown as replicate measurements from a single donor (n = 1), with three replicates per condition.

**Figure S9:**
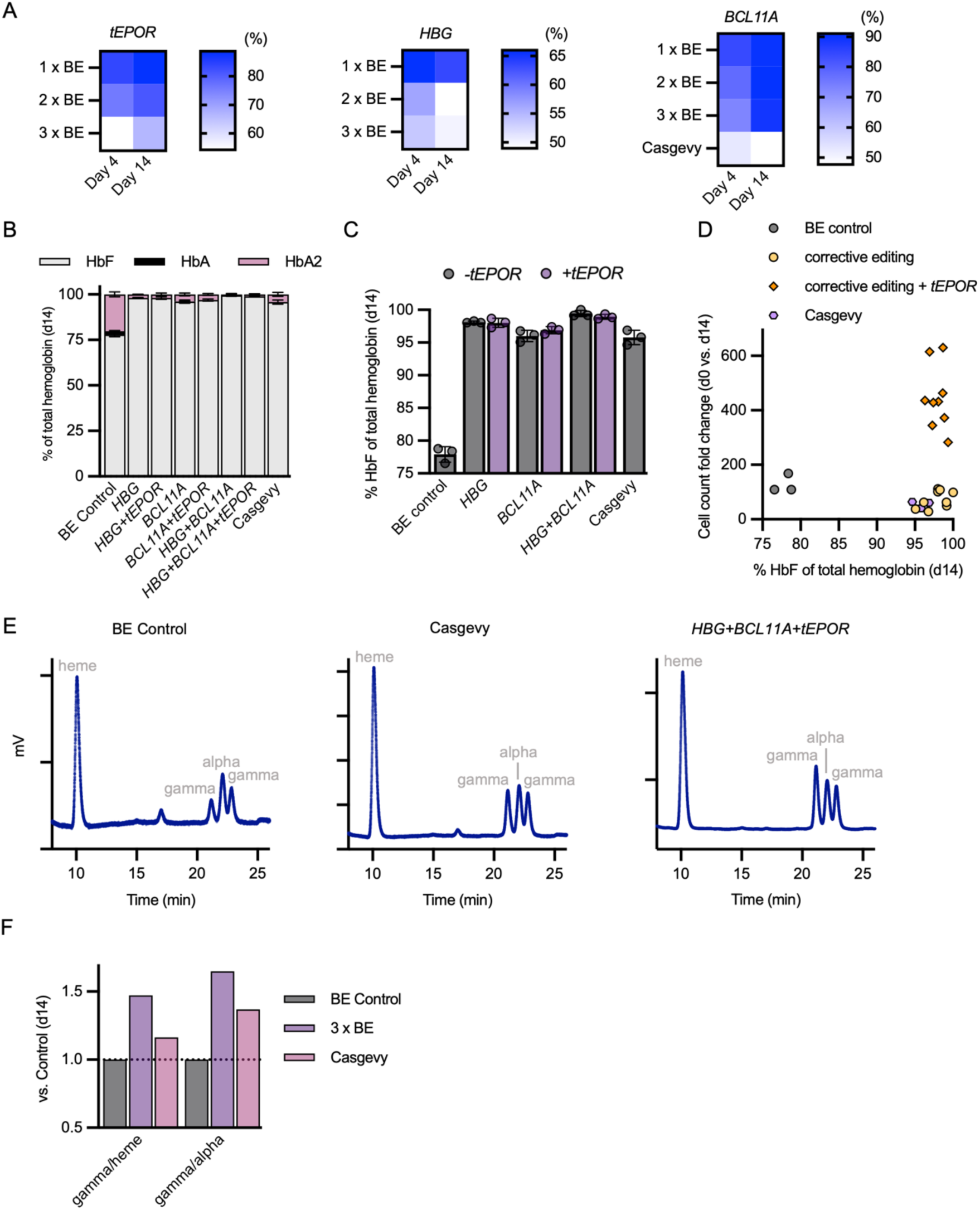
Multiplex base editing in β-thalassemia major HSPCs, related to Figure 5. **A.** Heatmaps showing editing frequencies at Day 4 and Day 14 of erythroid differentiation. **B-C.** Percentage of HbF among total hemoglobin following erythroid differentiation, determined by hemoglobin tetramer HPLC. **D**. Correlation between cell count fold change and HbF production at end of erythroid differentiation. **E.** Globin chains HPLC profiles obtained from the indicated cell lysates. Position of heme, alpha and of the two gamma peaks are indicated. **F.** Quantification of gamma amount relative to heme (gamma/heme) and to alpha (gamma/alpha) derived from HPLC measurements. Columns represent mean ± SD, and points represent replicate measurements from a single donor (n = 1; 3 replicates per condition).

**Figure S10:**
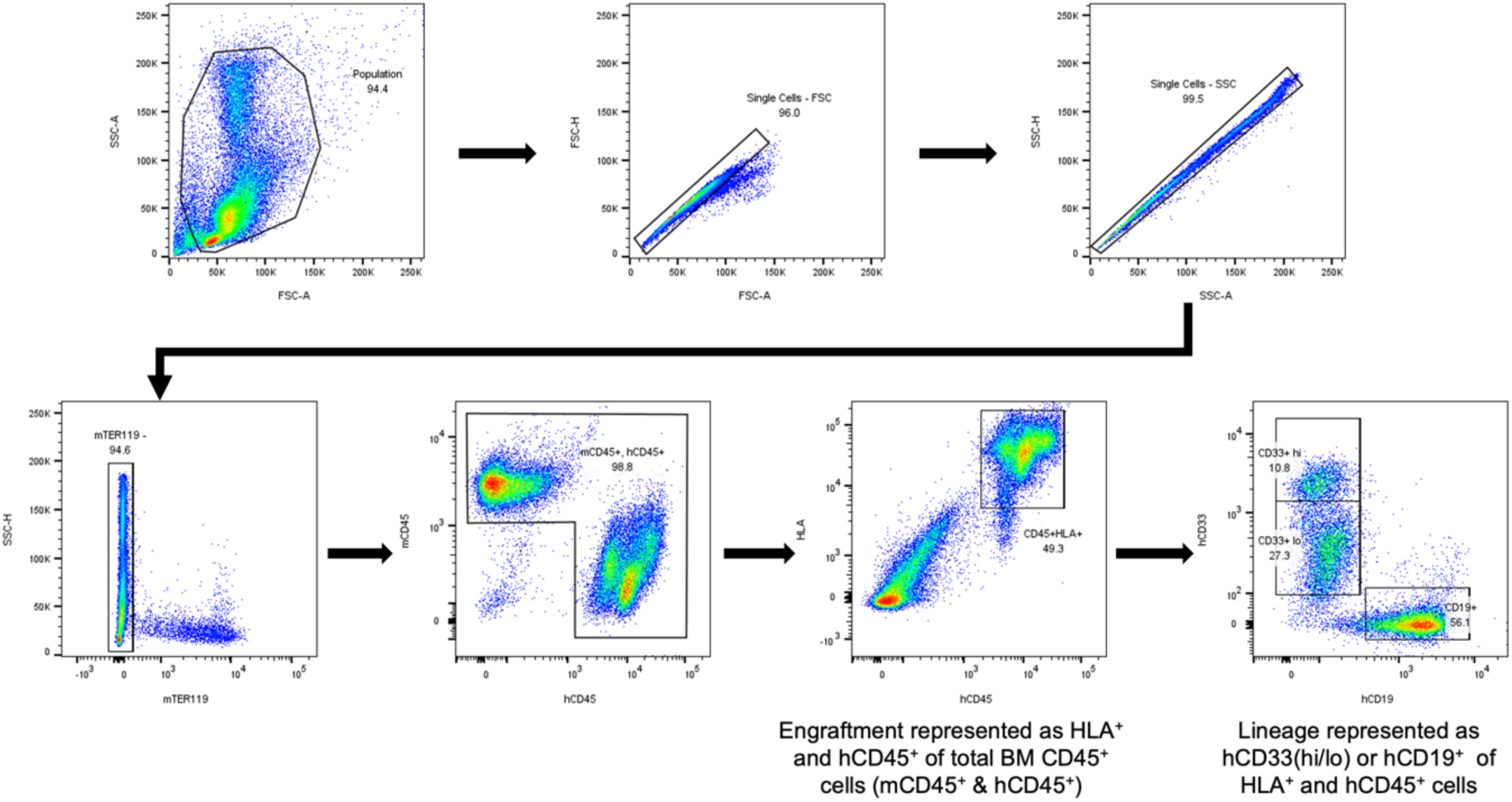
Engraftment and lineage gating scheme, related to Figure 6. Representative flow cytometry gating strategy for analysis of human HSPC engraftment in xenografted mouse bone marrow. Cells were first gated to exclude debris and doublets, followed by selection of live cells. Human engraftment was defined as hCD45^+^ and HLA^+^ cells within total bone marrow CD45^+^ cells (including mCD45^+^ and hCD45^+^ populations). Lineage distribution was determined within the hCD45^+^/HLA^+^ population, with myeloid cells defined as hCD33^+^ (including CD33hi/lo subsets) and B cells defined as hCD19^+^ cells.

**Figure S11:**
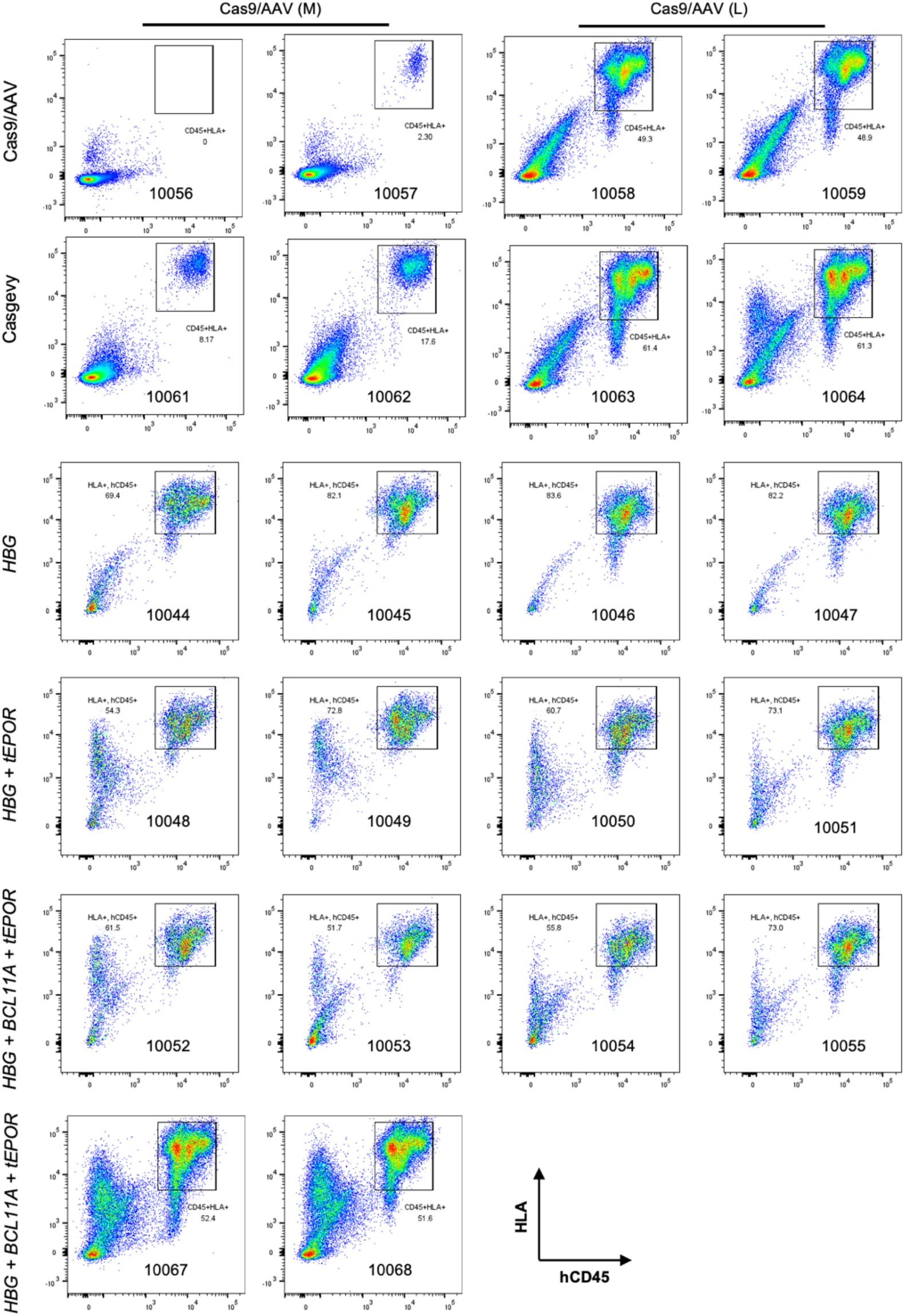
Flow cytometry analysis of human HSPC engraftment in xenografted mouse bone marrow, related to Figure 6. Bone marrow was harvested at 16 weeks post-injection of human HSPCs into NBSGW mice. Antibody staining and flow cytometry was used to quantify human engraftment (defined as the percentage of hCD45⁺/HLA⁺ leukocytes within the total CD45⁺ bone marrow compartment (mouse + human)). Flow cytometry plots for each animal are shown.

**Figure S12:**
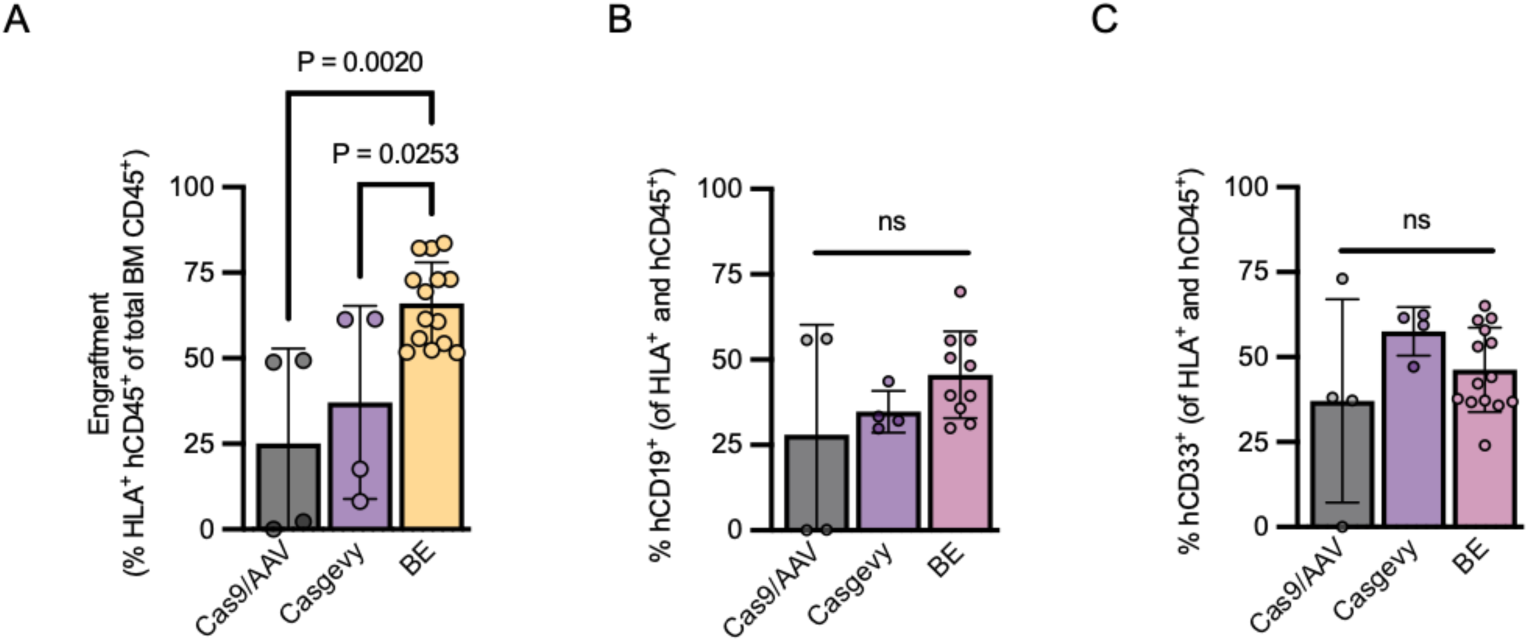
Engraftment and lineage distribution across combined BE groups (1x, 2x, 3x), related to Figure 6. **A.** Human hematopoietic engraftment in the mouse bone marrow at 16 weeks post-transplantation, shown as the percentage of HLA⁺ and hCD45⁺ cells within total bone marrow CD45⁺ cells. **B.** Percentage of human lymphoid (CD19⁺) cells among the engrafted human hematopoietic population (HLA⁺ and hCD45^+^) at 16 weeks post-transplantation. **C.** Percentage of human myeloid (CD33⁺) cells among the engrafted human hematopoietic population (HLA⁺ and hCD45^+^) at 16 weeks post-transplantation. BE conditions (1x, 2x, and 3x) are pooled and shown as a single group. Cas9/AAV conditions are grouped together by medium and low AAV doses. Each dot represents one mouse (n = 22); bars show mean ± SD. Statistical significance was determined using one-way ANOVA with Dunnett’s multiple comparisons test; ns, not significant (P > 0.05).

**Figure S13:**
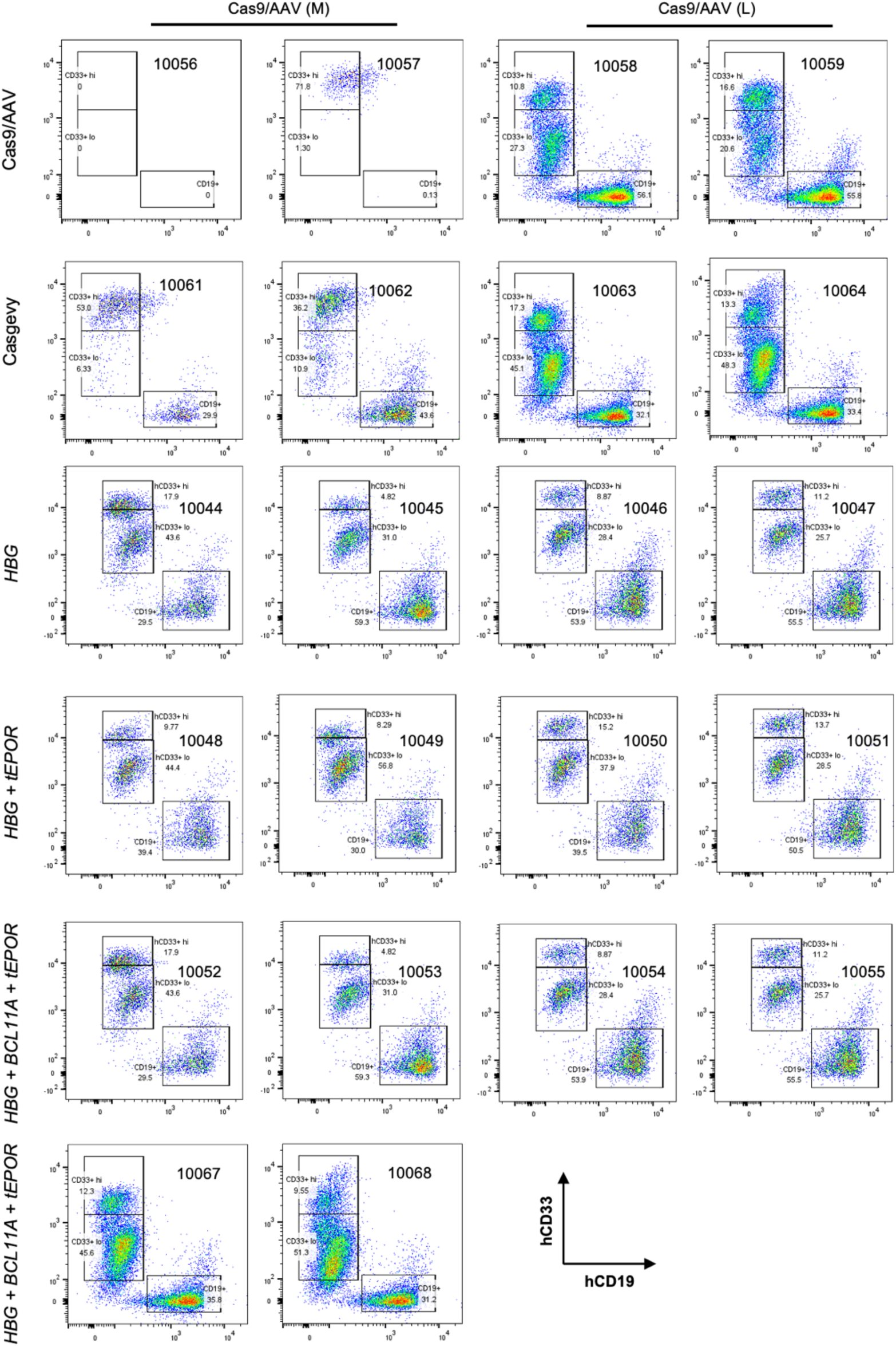
Flow cytometry analysis of human lineage reconstitution in xenografted mouse bone marrow, related to Figure 6. Bone marrow was harvested at 16 weeks post-injection of human HSPCs into NBSGW mice. Antibody staining and flow cytometry was used to quantify human myeloid and lymphoid reconstitution within the human graft. CD33⁺ subsets were further divided into CD33^hi^ and CD33^lo^ fractions based on expression intensity, representing more mature vs. less mature myeloid populations, respectively. Flow cytometry plots for each animal are shown.

**Figure S14:**
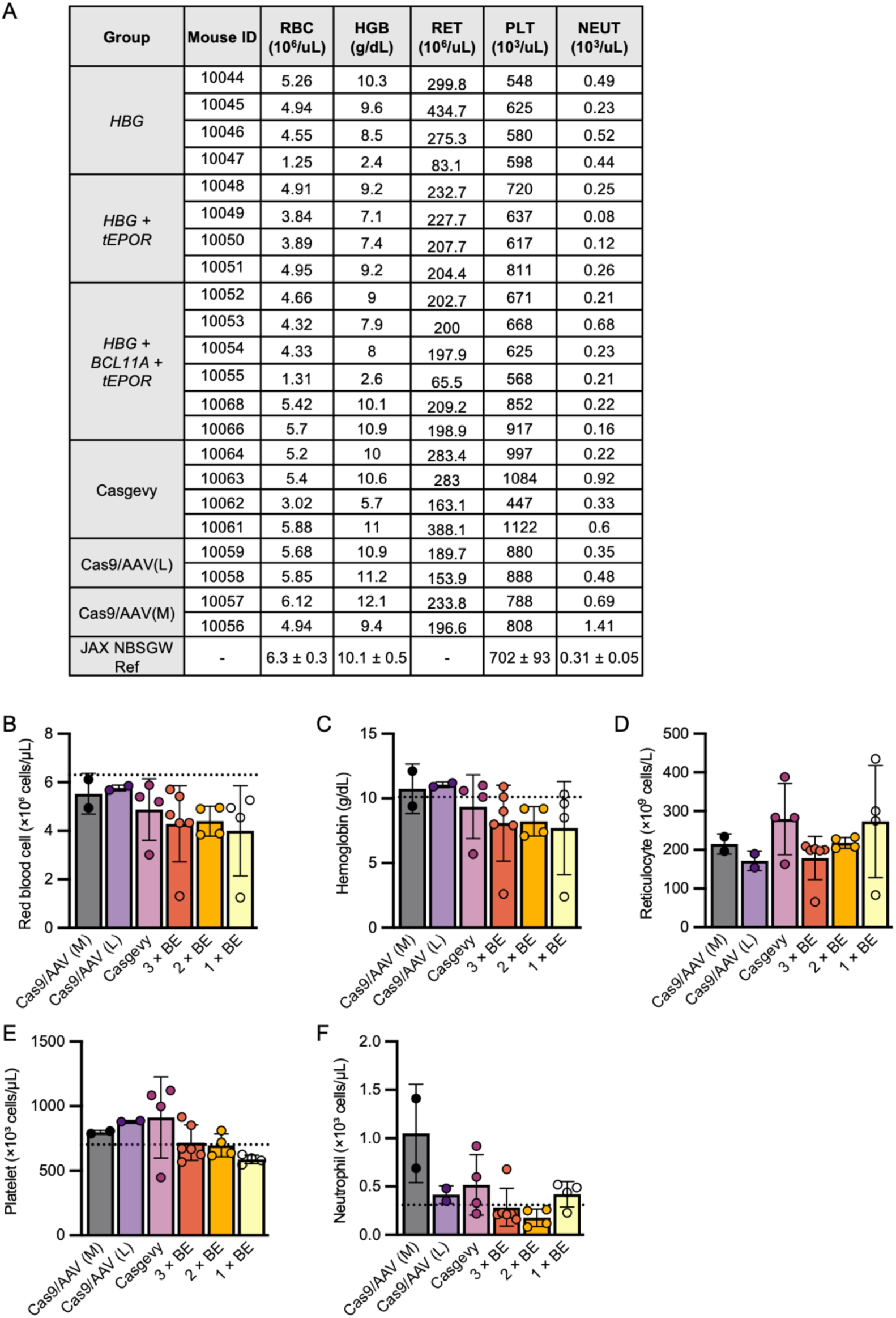
Peripheral blood hematologic profiling in xenografted mice, related to Figure 6. **A.** Sysmex complete blood count (CBC) measurements from NBSGW mice 16 weeks after transplantation with human HSPCs. Quantified parameters include red blood cells (RBC), hemoglobin (HGB), absolute reticulocytes (RET), platelets (PLT) and neutrophils (NEUT). Reference values for age-matched NBSGW mice are shown for comparison. **(B-F)** Quantification of individual hematologic indices across treatment groups. Dotted horizontal lines indicate the expected average values for age-matched NBSGW mice.

## STAR*METHODS

Detailed methods are provided in this paper and include the following:

1. KEY RESOURCES TABLE
2. RESOURCE AVAILABILITY

1. Lead contact
2. Materials availability
3. Code availability
3. EXPERIMENTAL MODEL AND SUBJECT DETAILS

1. Patient and healthy donor cells
2. Animal ethics statement
4. METHOD DETAILS

1. Plasmid construction
2. HeLa cell culture and transfection
3. *In vitro* transcription
4. *In vitro* culture of HSPCs
5. Genome editing of HSPCs
6. In vitro erythroid differentiation
7. Cell count and viability
8. Extraction and PCR amplification of genomic DNA
9. Sanger sequencing analysis of editing frequency
10. Targeted amplicon sequencing analysis of editing frequency
11. Digital PCR analysis of editing frequency
12. Immunophenotyping of differentiated erythroid cells
13. gRNA off-target analysis
14. Colony-forming unit (CFU) assay
15. Hemoglobin tetramer analysis
16. Single globin chain analysis
17. Xenotransplantation into NBSGW mice
18. In vivo engraftment & lineage analysis
19. Peripheral blood analysis
20. Mixed-effects modeling for HbF and cell count
21. Statistical analyses

## STAR*METHODS

### KEY RESOURCES TABLE

| REAGENT or RESOURCE | SOURCE | IDENTIFIER |
| --- | --- | --- |
| Bacterial and virus strains |  |  |
| *E. coli* Competent Cells | New England Biolabs | Cat# C3040 |
| Antibodies |  |  |
| hCD45 V450 | BD Biosciences | Cat# 560368 |
| CD34 APC | BioLegend | Cat# 343510 |
| CD71 PE-Cy7 | Affymetrix | Cat# 25-0719-42 |
| CD71 V650 | BioLegend | Cat# 334122 |
| CD235a PE (GPA) | BD Biosciences | Cat# 555570 |
| Ghost Dye Red 780 | Tonbo Biosciences | Cat# 13-0865 |
| mTER119 BUV395 | BD Biosciences | Cat# 563827 |
| mCD45 PerCP/Cy5 | BioLegend | Cat# 103132 |
| hCD45 AF488 | BioLegend | Cat# 304017 |
| HLA APC | BD Biosciences | Cat# 555555 |
| hCD19 PE | BD Biosciences | Cat# 555413 |
| hCD33 BV421 | BD Biosciences | Cat# 562854 |
| GPA/hCD235a PE | BioLegend | Cat# 349106 |
| CD71 APC | BD Biosciences | Cat# 551374 |
| Band3/CD233 FITC | Miltenyi Biotec | Cat# 130-119-780 |
| Biological samples |  |  |
| Healthy donor HSPCs (CD34-enriched) | Fred Hutchinson Cancer Center Hematology Core | N/A |
| Sickle cell disease HSPC (CD34-enriched) | Vivien Sheehan Laboratory at Emory University | N/A |
| β-thalassemia HSPCs (CD34-enriched) | UCSF | N/A |
| Chemicals, peptides, and recombinant proteins |  |  |
| DMEM | Hyclone | Cat# SH30022.01 |
| SFEM II | Stemcell | Cat# 09605 |
| MethoCult H4435 enriched methylcellulose medium | STEMCELL | Cat# 04435 |
| Penicillin-streptomycin | Invitrogen | Cat# 15140122 |
| Puromycin | Sigma-Aldrich | Cat# P8833 |
| Recombinant human stem cell factor (SCF) | Peprtech | Cat# 300-07 |
| Recombinant human Flt3-ligand (Flt3-L) | Peprtech | Cat# 300-19 |
| Recombinant human (IL-3) | Peprtech | Cat# 200-03 |
| Recombinant human interleukin-6 (IL-6) | Peprtech | Cat# 200-06 |
| Recombinant human thrombopoietin (TPO) | Peprtech | Cat# 300-18 |
| Recombinant human Erythropoietin (EPO) | Peprtech | Cat# 100-64 |
| Recombinant human transferrin | Sigma-Aldrich | Cat# T0665 |
| Human AB serum | Atlanta Biologicals | Cat# S40110 |
| Human plasma | Umbilical cord blood | N/A |
| Insulin | Sigma-Aldrich | Cat# I9278 |
| Heparin | Sigma-Aldrich | Cat# H3149 |
| Trypan blue | Thermo Fisher | Cat# 15250-061 |
| QuickExtract DNA Extraction Solution | Lucigen | Cat# QE09050 |
| Phusion Green Hot Start II High-Fidelity PCR Master Mix | Thermo Fisher Scientific | Cat# F566 |
| N1-methylpseudouridine-5'-triphosphate | TriLink | Cat# N-1081 |
| CleanCap AG | TriLink | Cat# N-7113 |
| CRISPR-Cas9 Protein, Streptococcus pyogenes, Recombinant | Aldevron | Cat# 9212 |
| Critical commercial assays |  |  |
| LIVE/DEAD Fixable Green Dead Cell Stain Kit | Invitrogen | Cat# L23101 |
| EasySep Human CD34 Positive Selection Kit II | STEMCELL | Cat# 17856 |
| Monarch RNA Cleanup kit | New England Biolabs | Cat# T2040 |
| Murine RNase inhibitor | New England Biolabs | Cat# M0314 |
| Lipofectamine 2000 | Thermo Fisher Scientific | Cat# 11668019 |
| P3 Primary Cell 4D-Nucleofector X Kit S | Lonza | Cat# V4XP-3024 |
| Monarch Genomic DNA Purification Kit | New England Biolabs | Cat# T3010 |
| HiScribe T7 High Yield RNA Synthesis Kit | New England Biolabs | Cat# E2040 |
| HiScribe T7 mRNA Kit with CleanCap AG | New England Biolabs | Cat# E2080 |
| QIAcuity Probe PCR Kit | Qiagen | Cat# 204443 |
| experimental models: Cell lines |  |  |
| Hela | Jia Liu laboratory | N/A |
| Software and algorithms |  |  |
| FlowJo (v10.7.1) | BD Biosciences | N/A |
| GraphPad Prism (v9.0) | GraphPad | N/A |
| TotalChrom | PerkinElmer | N/A |
| R (v4.3.1) | R Foundation for Statistical Computing | N/A |
| lme4 (v1.1-37) | R package | N/A |
| emmeans (v1.11.1) | R package | N/A |
| Cutadapt (v5.1) | Developed by Marcel Martin | https://cutadapt.readthedocs.io |
| FastQC (v0.12.1) | Babraham Bioinformatics | https://www.bioinformatics.babraham.ac.uk/projects/fastqc |
| MultiQC (v1.29) | Developed by Philip Ewels | https://multiqc.info |
| CRISPRessoBatch (v2.3.2) | Developed by Luca Pinello | https://crispresso.pinellolab.partners.org |
| Other |  |  |
| Lonza 4D-Nucleofector | Lonza | N/A |
| FACSAria II | BD Biosciences | N/A |
| CellDrop | DeNovix | N/A |
| Countess cell counter | Bio-Rad | N/A |
| QIAcuity dPCR system | Qiagen | N/A |

## RESOURCE AVAILABILITY

### Lead contact

Further information and requests for resources and reagents should be directed to and will be fulfilled by the lead contact, Kyle Cromer.

### Materials availability

All unique reagents generated in this study are available from the lead contact with a completed Materials Transfer Agreement.

### Code availability

All custom scripts used for statistical analysis are available from the Lead Contact upon reasonable request. No custom code was generated for NGS data processing. Any additional information required to reanalyze the data reported in this paper is available from the lead contact upon request.

## EXPERIMENTAL MODEL AND SUBJECT DETAILS

### Patient and healthy donor cells

Healthy donor G-CSF-mobilized, CD34-enriched peripheral blood HSPCs were purchased from Fred Hutchinson Cancer Center Hematology Core. Sickle cell disease CD34^+^ HSPCs were provided by the Sheehan laboratory at Emory University (IRB: 00034535). β-thalassemia samples were obtained as peripheral blood mononuclear cells (PBMCs) (IRB: 21-33506) and CD34-enriched using the EasySep Human CD34 Positive Selection Kit II according to the manufacturer’s protocol (STEMCELL Technologies).

### Animal ethics statement

All animal procedures were performed in accordance with the guidelines provided by the Emory University Institutional Animal Care and Use Committee (IACUC). The study was reviewed and approved under protocol number 202000101. All efforts were made to minimize animal suffering and to reduce the number of animals used. Mice were housed in a pathogen-free facility with a 12-hour light and dark cycle and provided ad libitum access to food and water. Euthanasia was performed via anesthetization with isoflurane by drop-chamber until the loss of pedal reflex and subsequent cervical dislocation.

## METHOD DETAILS

### Plasmid construction

*EPOR* gRNAs were cloned into the pGL3-sgRNA expression vector carrying a U6 promoter and a puromycin resistance gene (Addgene, #51133) for subsequent transfection into HeLa cells. The genes encoding AncBE4max^25^ and CBE6^53^ were each cloned into a plasmid vector containing a T7 promoter, followed by the 5′ UTR, Kozak sequence, open reading frame (ORF), and 3′ UTR for subsequent *in vitro* transcription (IVT).

### HeLa cell culture and transfection

Hela cells (ATCC) maintained in Dulbecco’s Modified Eagle Medium (DMEM) (Hyclone) supplemented with 10% fetal bovine serum (FBS) and 1% penicillin-streptomycin at 37 °C with 5% CO_2_. Transfections were performed at approximately 70% density about 12 hours after seeding, 0.5 μg gRNA plasmids and 1 μg AncBE4max plasmids were transfected with 3 μL Lipofectamine 2000 (Thermo Fisher Scientific). The medium was replaced with fresh medium with puromycin (Sigma-Aldrich, 2 μg/mL) at 12 hours after transfection, cells were harvested at 3 days after transfection.

### *In vitro* transcription

The base editor (BE) plasmid was linearized by PCR amplification using transcription primers listed in **Supplementary Table 5**. *In vitro* transcription (IVT) was performed using the HiScribe T7 High Yield RNA Synthesis Kit or the HiScribe T7 mRNA Kit with CleanCap AG (New England Biolabs; E2040 or E2080). CleanCap AG (TriLink, N-7113) was incorporated during transcription where indicated. In some reactions, uridine triphosphate (UTP) was replaced with N1-methylpseudouridine-5′-triphosphate (TriLink, N-1081). To prevent RNA degradation, Murine RNase inhibitor (New England Biolabs) was included in IVT reactions. Transcription reactions were performed in 0.5x transcription buffer, with 4 mM CleanCap AG added where indicated. To optimize cap incorporation efficiency, different dinucleotide cap analog:GTP ratios were used, including 1:1 (∼50% capped), consistent with previously described conditions^54^, and 4:1 (∼80% capped). In some reactions using the HiScribe T7 mRNA Kit with CleanCap Reagent AG, transcription was performed in 0.5x transcription buffer supplemented with 4 mM CleanCap AG and 6 mM ATP (∼95% capped). Following transcription, the mRNA was purified using the Monarch Spin RNA Cleanup Kit (New England Biolabs) and eluted in either nuclease-free water (ddH₂O) or TE buffer (IDT). Purified mRNA was stored at -80 °C until use.

### *In vitro* culture of HSPCs

CD34^+^ HSPCs were cultured at 1e5 cells/mL in StemSpan Serum-Free Expansion Medium II (STEMCELL Technologies) supplemented with a four-cytokine cocktail (PeproTech), including 100 ng/mL stem cell factor (SCF), 100 ng/mL thrombopoietin (TPO), 100 ng/mL Fms-like tyrosine kinase 3 ligand (FLT3L), and 100 ng/mL interleukin-6 (IL-6), as well as 100 µg/mL streptomycin and 100 U/mL penicillin. Cells were maintained at 37 °C in a humidified incubator with 5% CO₂.

### Genome editing of HSPCs

Human CD34⁺ HSPCs were thawed 48 or 72 hours prior to electroporation and maintained in previously described cytokine-supplemented medium. Synthetic gRNAs incorporated 2′-O-methyl modifications at the first three and last three nucleotides and phosphorothioate linkages between the first three and last three nucleotides (Synthego). For mRNA-based editing, mRNA and gRNA were mixed at a 1:1 weight ratio prior to electroporation, with mRNA dosages varied as indicated in the manuscript; in selected conditions, 3% RNase inhibitor was added to the RNA mixture. For ribonucleoprotein editing, recombinant Cas9 protein (Aldevron) was complexed with gRNA for 15 minutes at room temperature prior to electroporation. Initial control RNP experiments were performed at a final concentration of 4 μM RNP with a 1:1.2 molar ratio of Cas9 to gRNA, whereas experiments designed to maximize editing efficiency were performed using 8 μM Cas9 with a 3.5-fold molar excess of gRNA as previously described^55^. Immediately prior to electroporation, cells were resuspended in either P3 buffer (Lonza) or B1mix buffer containing RNA or RNP complexes and electroporated using a Lonza 4D Nucleofector with programs DZ-100 or CM-137. The B1mix electroporation buffer consisted of 3.6 mM KCl, 10.8 mM MgCl₂, 62.5 mM Na₂HPO₄, 24.3 mM NaH₂PO₄, 10 mM sodium succinate, 18 mM mannitol, 2.8 mM myo-inositol, 0.88 mM Ca(NO₃)₂, 0.55 mM sodium pyruvate, 21.8 mM D-glucose, and 2.19% GlutaMAX, adjusted to pH 7.2, and was prepared fresh or stored at 4 °C for up to two weeks prior to use^59^. Following electroporation, cells were plated at 1e5 cells/mL in cytokine-supplemented medium and cultured under standard conditions for downstream analyses. For Cas9 and AAV editing conditions, Cas9 RNP electroporation was immediately followed by transduction with recombinant AAV6 donor vectors at a multiplicity of infection of 625 (low dose), 2,000 (medium dose), or 5,000 (high dose). Low dose Cas9/AAV condition was incubated for 24 hours post-electroporation with 0.5 μM AZD7648 supplement to increase targeted integration frequencies, even at lower AAV doses^33^. Recombinant AAV6 vectors were produced in HEK293T cells (ATCC) and purified using commercial kits (Takara) as previously described^27^, and vector genome titers were determined by digital PCR to quantify vector genome copy numbers as previously described^56^. The AAV used in low dose condition was produced and purified as a large-scale research-grade AAV6 preparation by Charles River Laboratories.

### *In vitro* erythroid differentiation

Following editing, HSPCs were first cultured in HSPC maintenance medium (described above) for 2-3 days. Cells were then transitioned to erythroid differentiation conditions and cultured for 14 days at 37 °C and 5% CO₂ in SFEM II medium (STEMCELL Technologies).The SFEM II base medium was supplemented with 100 U/mL penicillin-streptomycin, 10 ng/mL stem cell factor, 1 ng/mL interleukin-3 (PeproTech), 3 U/mL erythropoietin (eBiosciences), 200 μg/mL transferrin (Sigma-Aldrich), 3% antibody serum (heat-inactivated; Atlanta Biologicals, Flowery Branch, GA, USA), 2% human plasma (from umbilical cord blood), 10 μg/mL insulin (Sigma-Aldrich), and 3U/mL heparin (Sigma-Aldrich). Cells were seeded at a density of 1e5 cells/mL during the first phase of differentiation (days 0-7). At day 7, cells were transitioned to the above media without interleukin-3 and seeded at 1e5 cells/mL. At day 11, cells were transitioned to the above media without interleukin-3 and with transferrin increased to 1 mg/mL and were seeded at 1e6 cells/mL.

### Cell count and viability

Cell number and viability were assessed by trypan blue exclusion (1:1 dilution) using a CellDrop automated cell counter (DeNovix) or Countess cell counter (Bio-Rad).

### Extraction and PCR amplification of genomic DNA

Genomic DNA from HeLa cells and HSPCs was isolated using QuickExtract DNA Extraction Solution (Lucigen) according to the manufacturer’s instructions. Briefly, cells were lysed in 20 μL QuickExtract solution per 5e4 cells. Genomic DNA from mouse bone marrow samples was extracted using the Monarch Genomic DNA Purification Kit (New England Biolabs). PCR was performed using 100 ng genomic DNA and Phusion Green Hot Start II High-Fidelity PCR Master Mix (Thermo Fisher Scientific). PCR amplification was carried out using a touchdown cycling protocol consisting of 30 cycles of 98 °C for 10 seconds, annealing for 15 seconds with the temperature decreasing from 68 °C to 58 °C at a rate of -1 °C per cycle, and extension at 68 °C for 60 seconds.

### Sanger sequencing analysis of editing frequency

Genomic regions spanning the target sites were PCR-amplified and subjected to Sanger sequencing. Indel frequencies were quantified using TIDE (Tracking of Indels by DE composition)^57^ or ICE (Inference of CRISPR Edits)^58^, and base editing efficiencies were quantified using EditR^26^ based on chromatogram peak decomposition.

### Targeted amplicon sequencing analysis of editing frequency

PCR products were purified using Gel Extraction Kit (QIAGEN) before construction of NGS libraries. Non-specific sequences in the PCR products were eliminated by gel electrophoresis. PCR products with different barcodes were pooled together for deep sequencing using the Illumina MiSeq (2x250) platform at Azenta. Raw sequencing reads were demultiplexed using Cutadapt (v5.1) based on sample-specific barcode sequences, allowing an error rate of 0.1 and quality trimming with a cutoff of Q20. Read quality was assessed using FastQC (v0.12.1), and summary reports were generated with MultiQC (v1.29). Processed reads were analyzed using CRISPRessoBatch (v2.3.2). For base editor experiments, quantification was performed using the --base_editor_output option with a quantification window centered at −10 and a window size of 20. For Cas9 nuclease experiments, quantification was performed using a quantification window centered at −3 relative to the predicted cut site, with a window size of 30. Editing and indel frequencies for each target site were determined from CRISPRessoBatch outputs.

### Digital PCR analysis of editing frequency

DNA samples were digested with NotI-HF (New England Biolabs) in rCutSmart buffer at 37°C for 5-15 minutes prior to digital PCR (dPCR) analysis. dPCR reactions were prepared using the QIAcuity Probe PCR Kit (Qiagen) with target-specific primer-probe sets (IDT) and loaded onto QIAcuity Nanoplates according to the manufacturer’s instructions. Reactions were partitioned and amplified on the QIAcuity dPCR system (Qiagen) with the following cycling conditions: 95 °C for 2 minutes, followed by 40 cycles of 95 °C for 15 seconds and 60 °C for 30 seconds. Absolute copy numbers were determined using QIAcuity Software Suite based on positive partition counts.

### Immunophenotyping of differentiated erythroid cells

Differentiated erythroid cells were analyzed by flow cytometry for erythrocyte lineage-specific markers using a FACS Aria II (BD Biosciences) at the UCSF Flow Cytometry Core. Edited and non-edited cells were analyzed using the following antibodies: hCD45 V450 (HI30; BD Biosciences), CD34 APC (561; BioLegend), CD71 PE-Cy7 (OKT9; Affymetrix), CD71 V650 (CYIG4; BioLegend), and CD235a PE (GPA) (GA-R2; BD Biosciences). To measure viability, cells were also stained with Ghost Dye Red 780 (Tonbo Biosciences) or a green fluorescent reactive dye (Invitrogen).

### gRNA off-target analysis

Potential off-target sites for *EPOR* gRNAs were predicted using the COSMID webtool^29^. Candidate sites were selected based on sequence similarity and mismatch tolerance within the gRNA protospacer sequence and prioritized by a COSMID score threshold of ≤5.5 as defined by prior work to capture all validated off-target sites when performing *ex vivo* editing of primary human HSPCs using high-fidelity Cas9.

### Colony-forming unit (CFU) assay

Edited CD34⁺ HSPCs were plated in methylcellulose-based medium (MethoCult H4434, STEMCELL Technologies) supporting colony formation at a density of 500 cells per 35-mm dish. Cultures were incubated at 37 °C with 5% CO₂ for 14 days to allow colony formation. Colonies were scored and colony types were classified as CFU-GEMM, CFU-GM, BFU-E, or CFU-E based on morphological criteria. Individual colonies were picked under microscopic guidance using a sterile pipette tip. Each colony was transferred into lysis buffer, and genomic DNA was prepared for genotyping to determine editing outcomes at the target loci. For the *HBG*-only condition, individual colonies were analyzed at the *HBG* locus. For 3x BE conditions (*HBG + BCL11A + tEPOR*), each colony was analyzed at all three loci. For Casgevy-edited samples, genotyping was performed at the *BCL11A* locus only. Due to limited colony numbers in the Cas9/AAV condition, insufficient material was available for genotyping.

### Hemoglobin tetramer analysis

Frozen pellets containing approximately 1e6 *in vitro*-differentiated erythroid cells were thawed and lysed in 30 μL of 0.03% Triton X-100 (made in water) for 10 minutes with constant shaking on a thermomixer (Eppendorf). Lysates were clarified by centrifugation at 13,000 RPM for 10 minutes at 4 °C. Native hemoglobin tetramers were analyzed by cation-exchange high-performance liquid chromatography (HPLC)using a PolyCAT A column (35 x 4.6 mm, 3 μM, 1500 Å; PolyLC Inc.) on a Flexar HPLC system (PerkinElmer) at room temperature, with absorbance monitored at 415 nm. Mobile phase A consisted of 20 mM Bis-Tris and 2 mM KCN (pH 6.94), and mobile phase B consisted of 20 mM Bis-Tris, 2 mM KCN, and 200 mM NaCl (pH 6.55). Hemolysates were diluted in buffer A and 20 μL was injected onto the column with an initial condition of 8% buffer B. Hemoglobins were eluted at a flow rate of 2 mL/minutes using a gradient from 8% to 40% B over 6 minutes, followed by 40% to 100% B over 1.5 minutes, and re-equilibration to 8% B for 3.5 minutes. Peak areas were quantified using TotalChrom software (PerkinElmer), and raw values were exported to GraphPad Prism (v9) for visualization and downstream analysis.

### Single globin chain analysis

Frozen pellets of approximately 1e6 cells *in vitro*-differentiated erythrocytes were thawed and lysed in 30 μL hemolysate buffer (EDTA 5mM, KCN 0.07%, pH 7.2) for 10 minutes with constant shaking on a thermomixer (Eppendorf). Lysates were clarified by centrifugation at 13,000 RPM for 10 minutes at 4 °C. Analysis of globin chains was performed by reverse-phase PerkinElmer Flexar HPLC system with a Vydac 214TP C4 column (250 × 4.6mm^60^, 5 μM, 300Å; Avantor, Inc.) at 20°C and detection at 280 nm. Mobile phase A consisted of 10.5% methanol made in acetonitrile and mobile B of 0.5% trifluoroacetic acid in water adjusted at pH 2.9 with NaOH. Hemolysate was diluted in B prior injection at a flow rate of 1 mL/minute in 49% A for 2 minutes, followed by 2 minutes in 50% A, and a 20-minute gradient to 60% A. The column was then equilibrated to 49% A for 6 minutes. Quantification of the area under the curve of the peaks was performed with TotalChrom software and raw values were exported to GraphPad Prism (v.9) for plotting and further analysis.

### Xenotransplantation into NBSGW mice

Eight- or twelve-week-old NOD.Cg-*Kit^W-41J^ Tyr* ^+^ *Prkdc^scid^ Il2rg^tm1Wjl^*/ThomJ (NBSGW) mice were used for xenotransplantation experiments. Prior to transplantation, mice were preconditioned with 20 mg/kg busulfan, 10 mg/kg administered interperitoneally 48 hours and 24 hours before transplantation. Edited human CD34^+^ HSPCs were transplanted via tail vein injection using a 30-gauge insulin syringe. Each mouse received 0.5-1e6 edited cells. Mice were monitored every 1-2 weeks for body weight and overall health. Acidified water was supplemented to diet to help prevent infection of immuno-compromised NBSGW mice. All animals were sacrificed 16 weeks post-transplantation, and tissues including peripheral blood, bone marrow, and spleen were collected for downstream analyses.

### *In vivo* engraftment & lineage analysis

Bone marrow was harvested 16 weeks after transplantation to measure human engraftment and evaluate lineage composition. Bone marrow cells were isolated from femur and tibia via centrifugation, and mononuclear cells were separated by density gradient separation using Ficoll-Paque Plus (GE Healthcare). Engrafted human cells were quantified via FACSymphony A5 Flow Cytometer (BD Biosciences) using antibodies against hCD45-Alexa Fluor 488 (HI30; BioLegend), HLA-APC (G46-2.6; BD Biosciences), mCD45-PerCP/Cy5.5 (30-F11; BioLegend), and mTER119-BUV395 (TER119; BD Biosciences). Human lineage output was assessed using antibodies against hCD19-PE (lymphoid cells) (HIB19; BD Biosciences) and hCD33-BV421 (myeloid cells) (WM53; BD Biosciences). All flow cytometry engraftment and lineage analyses were performed in post-acquisition software, FlowJo (v10.0). Editing frequencies in engrafted human cells were quantified by targeted sequencing.

### Peripheral blood analysis

Peripheral blood from transplanted mice was analyzed using an XN-1000V Hematology Analyzer (Sysmex) to measure white blood cells, red blood cells, hemoglobin, platelets, reticulocytes, neutrophils, and lymphocytes.

### Mixed-effects modeling for HbF and cell count

For analyses of HbF and cell count across multiple donors, we applied linear mixed-effects models using lme4 (v1.1.37) with donor treated as a random effect to account for repeated measurements. Values were log2-transformed prior to modeling. Models were specified as:

> log_2_(value)~condition+(1|donor)

Estimated marginal means were computed using emmeans (v1.11.1). Comparisons of each condition to control (*CCR5* CBE edit) were performed using Dunnett-adjusted contrasts. Predefined paired contrasts comparing matched *+tEPOR* and *-tEPOR* conditions were additionally evaluated, with P values adjusted using the Holm method. Effect sizes are reported as fold changes obtained by back-transformation of model estimates.

### Statistical analyses

Unless otherwise noted, all columns and error bars depict mean ± standard deviation (SD). Statistical significance was determined using one-way analysis of variance (ANOVA) followed by Dunnett’s multiple-comparisons test for comparisons with a control group. For pairwise comparisons, unpaired two-tailed Welch’s t-tests were performed with Holm-Šidák correction for multiple comparisons (α = 0.05). For experiments involving samples derived from multiple donors, linear mixed-effects models were used with donor treated as a random effect. Statistical analyses were performed using GraphPad Prism (v9) and R (v4.3.1) where indicated. A *P* value < 0.05 was considered statistically significant.

